# Unveiling the ultrastructural landscape of native extracellular matrix via lift-out cryo-FIBSEM and cryo-ET

**DOI:** 10.1101/2023.09.25.559261

**Authors:** Bettina Zens, Florian Fäßler, Jesse M Hansen, Robert Hauschild, Julia Datler, Victor-Valentin Hodirnau, Vanessa Zheden, Jonna Alanko, Michael Sixt, Florian KM Schur

## Abstract

The extracellular matrix (ECM) is a highly hydrated, three-dimensional network composed of various macromolecules and signaling factors. It serves as a structural scaffold for cells and plays an essential role in the regulation of numerous cellular processes, including cell migration, adhesion, and proliferation. Despite its importance in metazoans, structural knowledge is rudimentary on how the components of the matrisome are secreted, remodeled, and interact with each other and with surrounding cells. Specifically, the exact molecular assembly of important ECM fibers, such as fibronectin fibrils, fibrillin microfibrils, or Collagen-VI filaments has remained enigmatic. This is largely due to methodological limitations in specimen preparation for conventional room-temperature electron microscopy (EM).

To overcome these limitations, we have developed a cell culture-based 3D-ECM platform compatible with sample thinning by cryo-lift out focused ion beam (FIB) milling and cryo-electron tomography (cryo-ET). Our workflow involves the implementation of cell-derived matrices (CDMs) grown on EM grids, resulting in a highly adaptable and versatile tool to closely mimic ECM environments. This allows us to visualize native ECM and its components for the first time in their fully hydrated, cellular context. Our data reveals an intricate network of ECM fibers and their positioning with respect to matrix-secreting cells. In addition to D-spaced collagen fibers, we visualize previously unresolved fibrous structures, and an amorphous matrix co-assembling in proximity to ECM fibers and delineating the boundary between ECM and empty extra-cellular space. Intra- and extracellular granules presumably represent assembly intermediates of the ECM. Our results add to the structural atlas of the ECM and provide novel insights into ECM secretion, assembly and maintenance.

## Introduction

The extracellular matrix (ECM) is an intricate three-dimensional assembly of macromolecules and signaling factors and acts as a physical scaffold for ECM-residing cells. It controls cellular activities, such as proliferation or migration, via its biochemical, biomechanical, and biophysical properties. These properties are tissue-specific, depending on the origin of ECM-producing cells (*1*, *2*). Aberrant ECM composition and remodeling contribute to disease progression and alterations in the ECM are associated with aging, cancer metastasis, and fibrosis (*1*, *3*, *4*).

The ECM consists mostly of two classes of macromolecules: fibrous proteins (FPs) and proteoglycans (PGs). FPs include different types of collagens (Col, with 28 types existing in humans)(*5*), fibronectins (FN), fibrillins or elastins. They assemble the ECM scaffold and present soluble growth factors to cells (*1*, *2*, *6*). PGs form complexes with FPs or glycosaminoglycans (such as hyaluronan). Extensive interconnections between FPs and PGs are required for ECM fiber assembly and maintenance (*7–10*).

While we have a thorough understanding of the molecular inventory of the ECM, also due to proteomic studies in different tissues (*11–13*), the structural landscape of ECM fibrils and their interactions remain uncharted. Current approaches for studying ECM structures are mostly limited to the visualization of chemically contrasted specimens, where the employed conventional room temperature (RT) electron microscopy (EM) techniques are destructive to the strongly hydrated ECM environment. Finer details and importantly the molecular assembly of ECM components into fibrils or other higher-order arrangements cannot easily be discerned from such fixed and dehydrated ECM preparations. Beyond certain types of D-spaced collagen assemblies with a defined repeat length of 67 nm (e.g. Col-I or II) (*14–16*), there is no unambiguous consensus on the exact molecular assembly of many of the other ECM fibers, such as FN fibrils, fibrillin microfibrils, or Col-VI filaments. For these fibers, studies employing different EM or super-resolution fluorescence microscopy approaches reported different fiber dimensions with varying diameters or repeat length (**Table S1**). For example, Col-VI was suggested to form a unique fiber assembly among the collagen superfamily, with a 105-112 nm beaded repeat as revealed by electron EM (*17–19*). In contrast, immunogold labeling cryo-scanning transmission electron tomography (cryo-STET) data measured a Col-VI repeat pattern of only 85 nm (*20*). The challenge in reconciling these different observations is due to the use of varying experimental modalities, the fact that some ECM fibers are indeed variable in the sizes they can grow into, and that most measurements have been conducted *in vitro* or under dehydrated conditions not representative of physiological ECM assembly. A case in point are collagen fibers which spontaneously assemble *in vitro*, but not *in vivo* where they require additional proteins (*21*). Furthermore, FN, fibrillins, and elastins assemble deformable fibers (*22*, *23*), able to adapt to the tension exerted by cells and tissue.

The intricate and complex interplay between ECM components can be best studied within a native environment, such as in the context of matrix-secreting cells. However, this has been exacerbated by technological limitations in cryo-electron tomography (cryo-ET) which enables visualizing specimens in 3D under virtually artifact-free conditions (*24*). Specifically, the thickness of ECM specimens, extending into the tens of micrometer range, requires additional sample thinning steps, by for example using cryo-focused ion beam scanning electron microscopy (FIBSEM). Cryo-FIBSEM has been successfully applied to a variety of specimens, i.e., when using bulk milling on isolated adherent cells (*25*, *26*) or even small organisms, such as *C. elegans* (*27*). The lift-out technique has been introduced to obtain lamellae of samples that are otherwise incompatible with conventional bulk milling approaches (*24*, *28*).

Another aggravating factor for the structural annotation of ECMs is the heterogeneity of tissue-derived ECM material. In contrast, cell derived matrices (CDMs) are a highly adaptable and versatile tool that is increasingly used to recapitulate the complexity of native tissue ECM (*29–31*). To obtain CDMs, ECM-producing cells such as fibroblasts are cultured over several weeks to produce a 3D matrix that closely resembles the tissue these cells originate from. Given their single-cell type origin, CDMs provide the advantage of high reproducibility, genetic tractability, and homogeneity. Hence, CDMs are used in fundamental research focusing on cell motility and cell proliferation, as well as tissue engineering and regenerative medicine (*29*, *32–35*).

Here, we have implemented CDMs in a cell culture-based 3D-ECM platform, which is compatible with sample thinning by cryo-lift out FIB milling and cryo-ET. The versatility and adaptability of CDMs enables us to closely mimic ECM environments and to study natively preserved ECM structures in 3D. Our cryo-electron tomograms of CDM reveal an intricate network of ECM fibers in context of matrix-secreting cells and provide further insights into the still open questions on the molecular assembly of ECM components into a functional and intricate 3D matrix.

## Results and Discussion

### On-grid CDM growth and characterization

We adapted a previously published protocol (*31*), to render CDMs produced by human telomerase immortalized foreskin fibroblasts (TIFFs) compatible with downstream cryo-liftout FIBSEM and cryo-ET experiments. Employing our recently published grid holders to facilitate long-term cell culture on EM-grids (*36*), we seeded TIFFs onto EM grids and performed time course studies to determine the optimal culture conditions and growth time to generate fully formed CDMs (**Figure 1A,B, see Materials and Methods**). Collagen secretion from cells could be already observed shortly after reaching cell confluency (designated as Day 0) and at Day 7 collagen fibers had formed. After 14 days, collagen assembly in CDMs reached an average height of 14.8 µm (n = 7; SD = ± 2.8 µm) when measured in confocal microscopy, which did not substantially increase when cultivated for longer time points. Mass spectrometry of day-14 CDMs identified 110 ECM proteins, supporting that at this time point a full matrix assembly had formed. Among the 25 most abundant matrisome components were FN-I, Col-I, Col-VI, the collagen crosslinker Tenascin and sugar-binding proteins such as Galectin-1 and Galectin-3 (**Table S2**). Correspondingly, FN fibers were also highly enriched in our Day 14 CDMs when visualized via immunofluorescence microscopy, which also clearly revealed a multilayer of cells embedded within ECM (**Figure 1C**).

**Figure 1:**
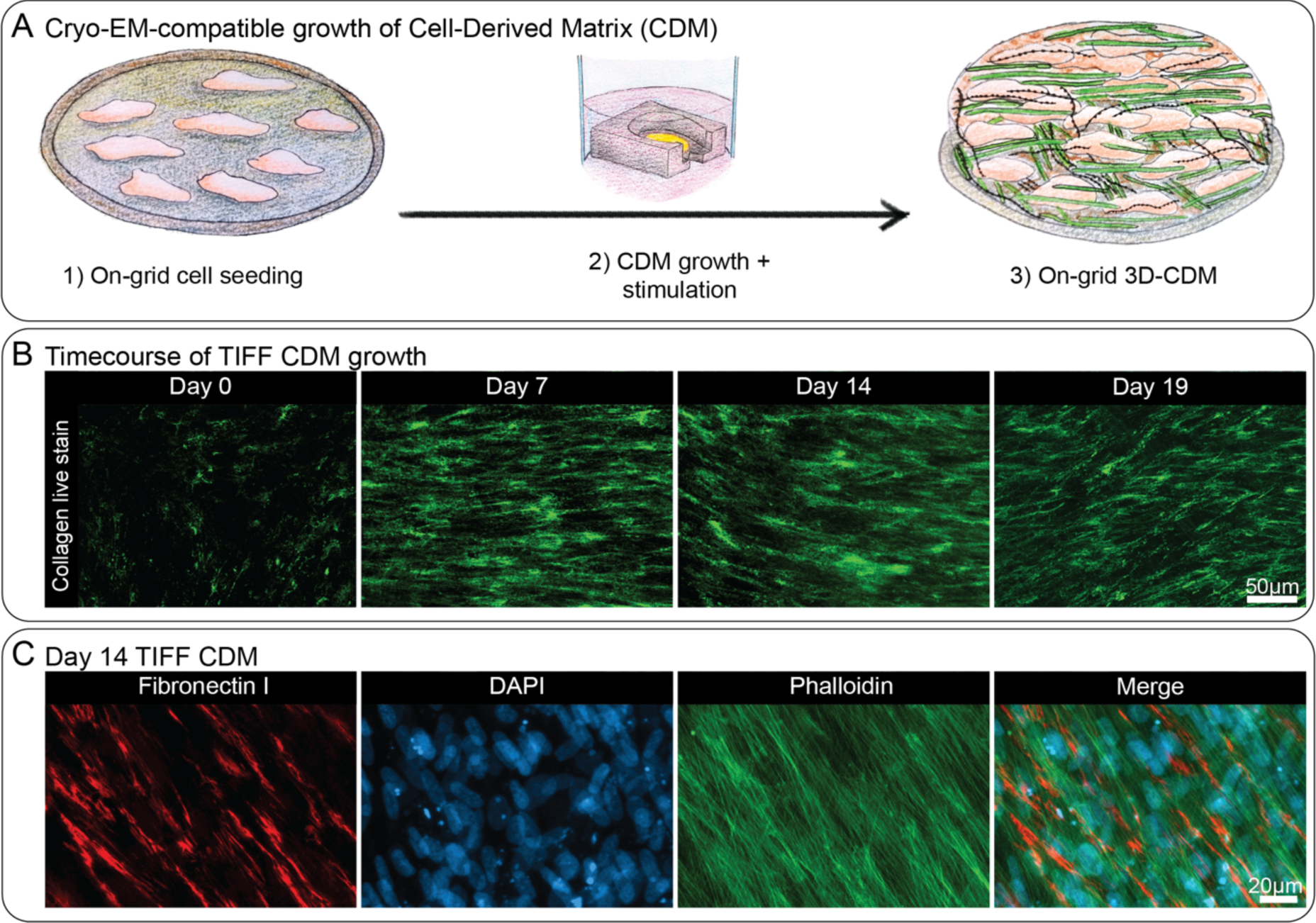
On-grid CDM generation and characterization. **A)** Schematic depiction of CDM growth on EM grids (see Materials and Methods for details). **B)** Time-course of on-grid collagen fiber deposition over 19 days. CDMs were live-stained with the collagen binding protein CNA35-EGFP and imaged by confocal microscopy. Shown here are maximum intensity Z-projections of exemplary CDM areas on different representative EM grids on Day 0 (start of ascorbic acid treatment after reaching cell confluency), Day 7, Day 14, and Day 19. **C)** TIFF CDMs were grown for 14 days, fixed, and stained with an anti-FN I antibody, and DAPI and phalloidin to visualize the nucleus and actin cytoskeleton, respectively. Specimens were imaged by confocal microscopy and an exemplary region of a CDM is shown here as maximum intensity Z-projection of each staining as well as a merge of all three stainings. Scale bar dimensions are shown in the figure.

To further characterize our CDMs we performed RT array tomography via scanning electron microscopy (SEM) of thin-sectioned CDMs (**Figure S1**). Array tomography sacrifices nativity and resolution due to the involved chemical fixation and dehydration process, but allows large-field volumetric imaging with improved resolution in the Z-axis compared to confocal imaging. Segmentation of the 3D array tomography data revealed cells to be adopting a flat and extended morphology stacked on top of each other, with ∼10 cells overlapping each other over a height of 15 μm. Cells were embedded in ECM, as judged by the presence of what we assumed to be collagen fibers (**Figure S1A,B**). Cell and ECM fibre orientation showed a clear dependence on the Z-height of the CDM (**Figure S1C**), where fibres aligned with the long axis of the cell, and changed dependent on their height positioning in the cell multilayer. Altogether, these results confirmed that CDMs harvested on or later than Day 14 represent *bona fide* ECM assemblies and allow visualization of ECM components in their native environment of matrix-secreting cells.

### Vitrification optimization and correlative imaging of CDMs

Due to their height and comparatively high free water content, CDMs exceed the vitrification potential of plunge freezing. Hence, to achieve optimal vitrification we performed high-pressure freezing (HPF) of on-grid CDMs (**Figure S2A**). Vitrification status after HPF can only be evaluated in cryo-transmission electron microscopy (cryo-TEM) via observation of ice crystal reflections in thinned lamellae. Hence, upon vitrification, we first imaged fluorescently-labelled CDMs via cryo-fluorescence light microscopy, to judge CDM and grid integrity and to define regions of interest (ROI) (**Figure S2B**). We then performed lift-out cryo-FIB milling to obtain CDM-containing lamellae that are thin enough to be subjected to cryo-TEM (**Figure S2C**). Initial vitrification trials without the use of a cryoprotectant resulted in lamellae showing high-contrast features in cryo-TEM, but also incomplete vitrification (**Figure S3A**). We therefore tested in total 12 different buffer compositions containing cryoprotectant for overall vitrification, as well as the additional background they introduced (**Table S3**). Many different cryoprotectant conditions commonly used in other experimental settings (*37–40*) resulted in insufficient vitrification in all tries (**Figure S3B**). Others resulted in such high background that cellular and ECM features could not be properly discerned (**Figure S3C**). A degassed cryoprotectant solution containing 10% Dextran in 0.1 M phosphate buffer (PB) showed the highest success and resulted in complete vitrification in several samples with low additional background (**Figure 2, Figure S4**). Hence, we proceeded with this cryoprotectant buffer for our further experiments.

**Figure 2:**
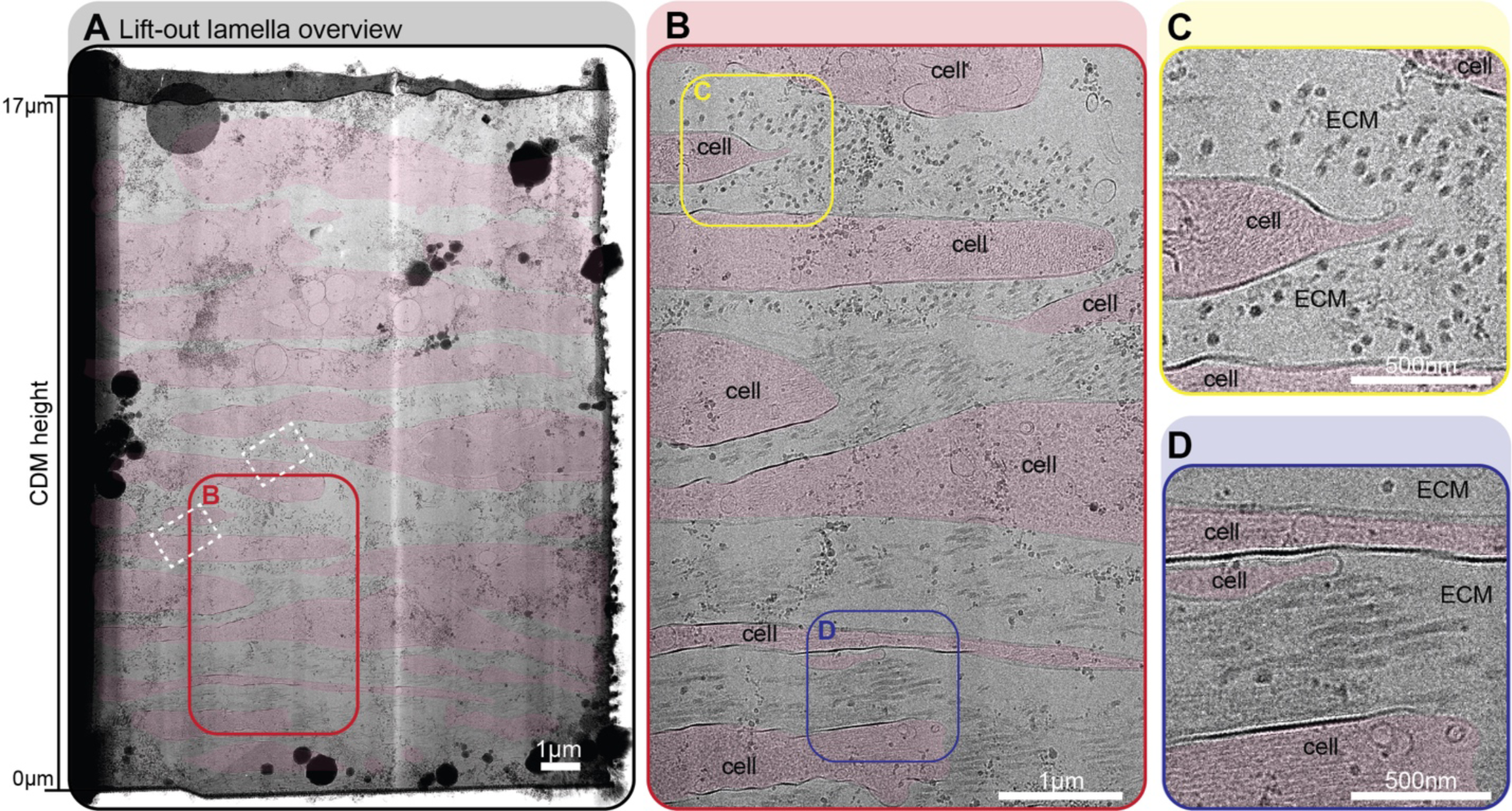
A fully vitrified cryo-lift out lamella reveals high-resolution ECM structures. **A-D)** Fully vitrified cryo-lift out lamella from a TIFF CDM grown for 16 days shown at different magnifications (cryoprotectant: 10% Dextran in degassed 0.1 M PB). Cell areas are annotated with transparent red color. **A)** Complete overview of the cryo-lift out lamella. The lamella covers roughly 17 µm of CDM depth, ranging from proximal to the EM grid substrate (Z = 0 µm) to close to the CDM surface (Z = 17 µm). White dashed rectangles denote the areas of acquisition for the tomograms shown in Figure 3. **B)** Zoom-in of the lamella as annotated with a red rectangle in (A). Two ROIs, highlighted by colored rectangles are shown at higher magnification in (C) and (D). **C)** Zoom-in into the CDM where ECM fibers are running perpendicular to the lamella, resulting in a cross-section view of fibers. **D)** Zoom-in into the CDM where ECM fibers are orientated parallel to the lamella, resulting in a side-view of fibers. Scale bar dimensions are annotated in the figure.

### Architecture of natively preserved CDMs

Cryo-TEM of our fully vitrified lift-out lamellae allowed us to visualize cell-ECM assemblies in a vertical cross-section view spanning over almost the entire depth of the CDM (**Figure 2**, and **Figure S4** for additional examples). The CDM was composed of several layers of cells, between which extracellular space was filled with ECM components (**Figure 2B**), in line with our observations made by array tomography. Most prominently we could observe thick ECM filaments running in orthogonal or parallel direction with respect to the plane of the lamella, again depending on their position with respect to CDM height. Empty areas in extracellular space devoid of any structures were also observed. From the bottom towards the top of the CDM the density of ECM fibres seemingly decreased, indicating a potential link of growth of new CDM preferentially on top with the expanding cell layers.

### Molecular view of natively preserved CDM

We acquired cryo-electron tomograms (n = 43, average thickness = ∼194 nm, standard deviation = ± 31.5 nm) from our lift-out lamellae, providing a high-resolution 3D view of the molecular components and the connections between cells and assembled ECM. Our tomograms revealed numerous cellular organelles or membranous compartments, besides hitherto undescribed ECM structures (**Figure 3, Figure S5, Movies S1-S2**).

**Figure 3:**
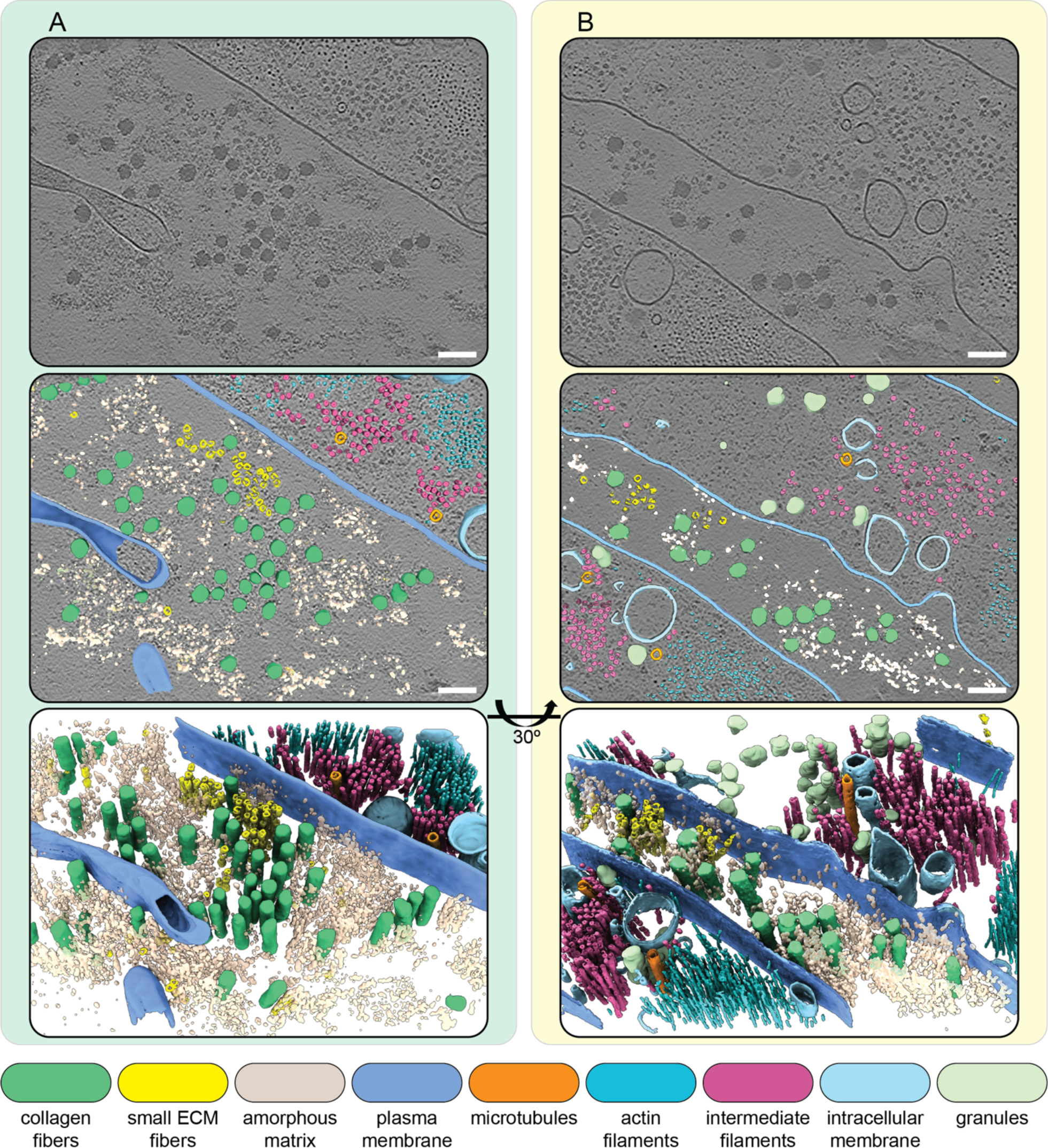
Molecular landscape of CDMs. **A, B)** Segmentations of two exemplary IsoNet-processed tomograms acquired on the cryo-lift out lamella shown in Figure 2. The top panels show a single central slice (1.71 nm thickness) of each respective tomogram. Cell and ECM fibers are aligned perpendicular to the lamella, resulting in a cross-section view of intra- and extracellular filaments. Middle panels show segmentations of tomograms overlaid over the tomogram slice. Bottom panels show an oblique view of just the segmentation volume. Scale bars indicate 100 nm. The color scheme for the different segmented cellular and ECM components is described in the figure.

Within cells we observed cytoskeletal filaments, easily identified via their characteristic diameters and appearances as intermediate filaments, actin filaments, and microtubules (**Figure 3A-B**, **Figure 4**). Some microtubules showed globular intraluminal microtubule-associated proteins, while other microtubules contained continuous filamentous densities, closely resembling structures of luminal actin recently observed in HAP60 and Drosophila S2 cells (*41*, *42*). In line with the orientation of ECM fibers along the long axis of the cell, also cytoskeletal filaments displayed a parallel orientation with respect to ECM fibers.

**Figure 4:**
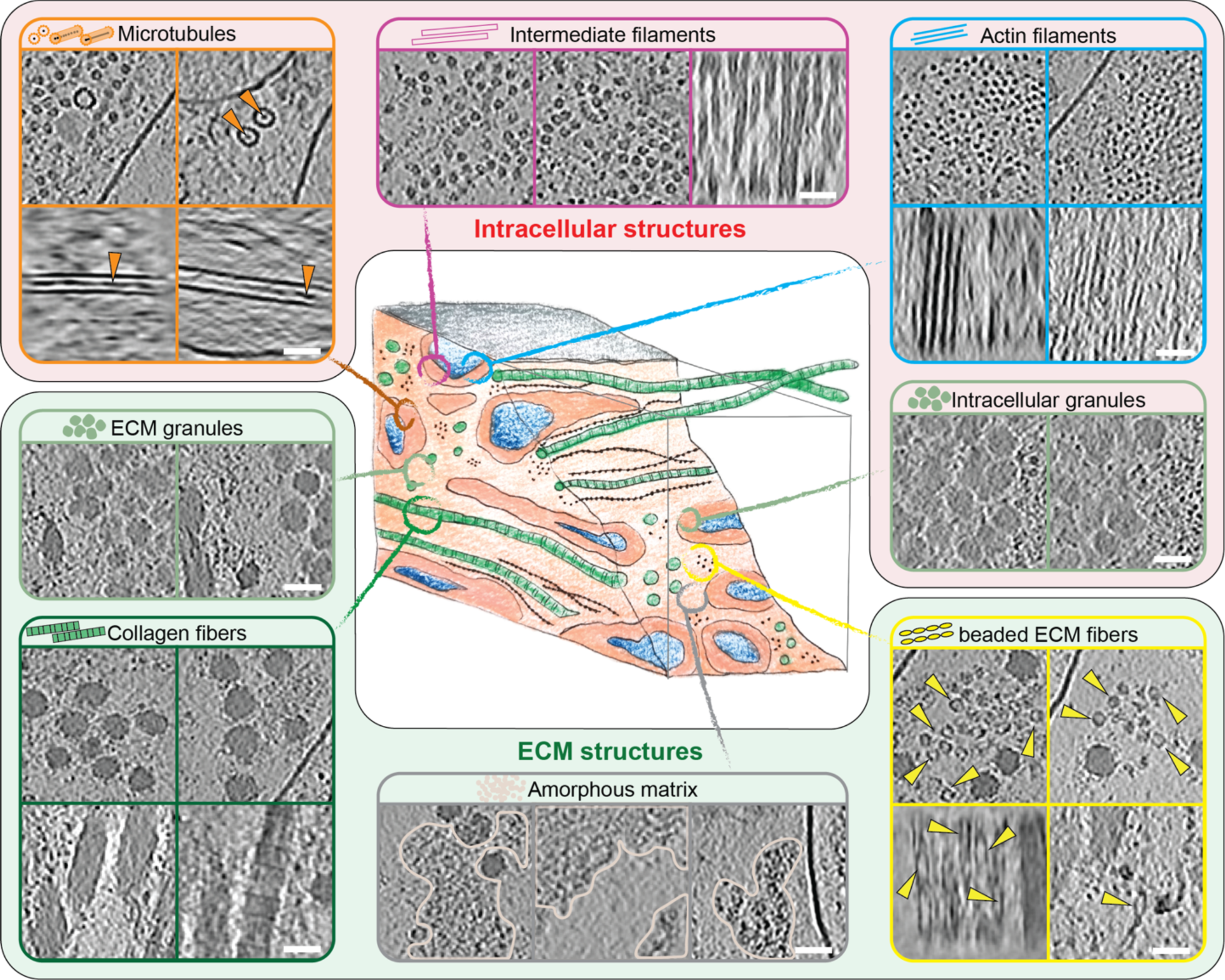
Gallery of extra- and intracellular structures in CDMs. Details of extra- and intracellular structures found in CDM tomograms. On the top intracellular features are shown and the bottom area displays ECM features. Each image is derived from an IsoNet-processed tomogram from a single z-slice (1.71 nm thickness). The scale bars indicate 50 nm.

Other well-resolved cellular features included endocytic sites with proteins assembling at the vesicular neck, and fully formed clathrin-coated vesicles, as judged by their dimensions and the clear presence of a protein coat (**Figure S6A-C**). We also observed numerous structures resembling endoplasmic reticulum (ER) compartments (**Figure S6D**) with areas of high local membrane curvature and other membrane-enclosed spherical entities.

### Distinct structural entities within the ECM

The ECM could be visually classified into four distinct structural entities (**Figure 4**). The most abundant and prominent ECM structures were large filaments with a diameter between 25-60 nm (**Figure 3, 4, S5, Movies S1-S2**), which we identified as collagen fibers, based on the banding pattern with a periodic D-spacing of ∼67 nm (**Figure S7A-B**). Based on their abundance in our cryo-ET data and our proteomic characterization of CDMs (**Table S2**) we assume these to be Col-I fibers. Our resolution does not reveal the triple helix arrangement of collagen, but shows them as highly-electron dense (and to a certain extent also more electron-dose sensitive) fibers, tightly embedded in other ECM components.

The second entity consisted of smaller, less regular filament assemblies distributed within the ECM (**Figure 3, 4**). These filaments had a diameter of ∼15.7 nm at their widest position (n = 110; SD = ± 1.3 nm) and a bead-like appearance with a node-to-node distance of ∼60.3 nm (n = 54; SD = ± 7.8 nm) (**Figure S7C-D**). The variation in both diameter and repeat pattern along the filament z-axis suggests extensibility and assembly variability. To our knowledge no similar ECM filament assembly has been yet visualized.

The third structural entity was an amorphous matrix that occupied large areas between filaments (**Figure 3 & 4, S8**). Accordingly, PGs were described to form a gel-like ground substance. In all cases the amorphous matrix was observed only in close proximity to ECM fibers, often sharply delineating the area filled with ECM components to empty extracellular space. This suggests a co-assembly between the components in these entities. Accordingly, it has been previously reported that coordinated presence and interactions between FP and PGs are required for proper ECM function since disruption or loss of PGs such as decorin leads to abnormal collagen assembly (*7–10*, *21*, *43–46*).

The last distinct component of the ECM we found were amorphous granules which appeared in clusters or sometimes also as isolated objects (**Figure S9A, Movie S3**). Granules of identical morphology were also often found within cells (**Figure 3B, S9B-C**). Here they again were either isolated or forming areas largely excluding other cellular components. In a few instances, release of granules into the extracellular space could also be observed (**Figure S9D**), leading us to speculate that these granules could represent ECM assembly intermediates. Accordingly, Col-VI has been described to first form large intracellular aggregates before secretion and eventually forming beaded fibers (*47*). However, the bead-like filaments we observed in our tomograms had dimensions inconsistent with previous Col-VI reports (**Table S1, Figure S7C-D**). Instead, fibrillin microfibrils in the resting state with a reported ∼56 nm periodicity and ∼10-20 nm diameter (**Table S1**) were closest to our measurements of the beaded filaments of ∼60 nm periodicity and ∼15 nm diameter (**Figure S7C-D**). Certain ECM filaments, such as Fibrillin microfibrils as well as FN fibers have different extensibility, providing adaptability of tissue. This can result in different measured periodicities depending on the tensile state, providing a potential explanation for the discrepancy between measurements among previous studies and also our data. Still, pending further experiments for the direct identification of the protein(s) constituting the beaded fibers in our data, their identity remains unclear.

### The absence of other filament-like structures in CDMs

Most strikingly, our data does not contain any other clearly discernible ECM fiber types, as one would expect based on our proteomics data or previous reports of the filamentous assemblies present within the ECM (such as FN fibers or Col-VI). However, within the amorphous matrix we regularly noticed features seemingly following a linear trajectory, but otherwise not standing out from the disordered matrix environment (**Movies S4 & S5**). This might suggest that certain ECM fiber assemblies might not take on a highly regular shape and morphology, but rather retain a structurally non-distinct shape that blends into the amorphous matrix, potentially due to decoration with PGs or other components. While this conclusion appears inconsistent with previous reports on the structure of Col-VI fibers (*47*), their apparent absence in our tomograms represents an interesting conundrum, deserving additional studies.

### Assigning identities to the unknown

We acknowledge that our analysis of the ECM is impeded by the major limitation and at the same time potential of any *in situ* cryo-ET study that performs exploratory structural biology, which is revealing all molecular components without discrimination. Unambiguous molecular assignments are only possible for structures that are already known (partially defeating the purpose of truly exploratory structural biology), when performing subtomogram averaging to determine protein identity from structure (*48*), or when using additional immunogold labeling strategies. As also revealed by our data, ECM filaments represent a challenging target for structure determination approaches due to their apparent variability. Future experiments using lift-out cryo-FIBSEM and cryo-ET, combined with novel image processing tools based on neural networks (for example (*49*)) or functional studies using genetic knockouts will be required to extend and annotate the gallery of ECM structures.

### Conclusions

Here, we present to our knowledge the first dedicated cryo-ET study of natively preserved ECM. Our workflow implementing CDMs as tools to study ECM assemblies allows visualizing hitherto undescribed structures and sets the stage for follow-up studies. This includes the structural and functional characterization of the molecular sociology of natively preserved ECMs. Specifically, based on the hypothesis that changes in matrisome composition of ECMs of different origin are reflected on a structural level, cryo-ET of CDMs now allows studying how ECM-specific topologies define tissue homeostasis and underlying cellular behavior. Hence, CDMs derived from cell types of different origin, e.g. skin, lung, mammary or cancer-associated fibroblasts (CAF), offer an appealing avenue for comparative analysis of ECMs to obtain a clearer depiction of the role of individual ECM components in defining specialized tissue-specific matrices. Specifically, this could be achieved via an integrative approach using a combination of molecular imaging *via* cryo-ET, proteomics analysis *via* mass spectrometry (MS) followed by functionally characterizing the role of specific matrix components using CRISPR-Cas9 knockout approaches. Together with pharmacological treatment to target specific ECM component this offers the possibility to manipulate ECM production.

## Acknowledgments

This research was supported by the Scientific Service Units (SSUs) of ISTA through resources provided by Scientific Computing (SciComp), the Lab Support Facility (LSF), and the Electron Microscopy Facility (EMF). Specifically, we thank Armel Nicolas and his team at the ISTA proteomics facility, and Alois Schloegl, Stefano Elefante and colleagues at the ISTA Scientific Computing facility for their support. We further thank Tommaso Constanzo and Ludek Lovicar at the Electron Microsocpy Facility (EMF), and Thomas Menner at the Miba Machine shop for their support. We also thank Wanda Kukulski (University of Bern) as well as Darío Porley, Andreas Thader, and other members of the Schur group for support and helpful discussions. Matt Swulius and Jessica Heebner provided great support in using Dragonfly. We thank Dorotea Fracciolla (Art&Science) for support in figure illustration.

We acknowledge funding support by the following sources: Austrian Science Fund (FWF) grant P33367 (to FKMS), the Federation of European Biochemical Societies (FEBS) (to FKMS), Niederösterreich (NÖ) Fonds (to BZ), FWF grant E435 (to JMH), European Research Council under the European Union’s Horizon 2020 research (grant agreement No. 724373) (to MS), and Jenny and Antti Wihuri Foundation (to JA). This publication has been made possible in part by CZI grant DAF2021-234754 and grant DOI https://doi.org/10.37921/812628ebpcwg from the Chan Zuckerberg Initiative DAF, an advised fund of Silicon Valley Community Foundation (funder DOI 10.13039/100014989) (to FKMS).

## Author contributions

Supervision and funding acquisition: M.S, F.K.M.S.; project administration: B.Z. and F.K.M.S.; conceptualization: B.Z. and F.K.M.S.; methodology: B.Z. and F.K.M.S.; investigation: B.Z., R.H., J.M.H, J.D., J.A., F.F., V.-V.H., and V.Z.; validation and formal analysis: B.Z., and F.K.M.S.; visualization: B.Z., J.M.H., J.D., R.H.; data curation: B.Z. and F.K.M.S.; writing—original draft: B.Z. and F.K.M.S.; writing—review and editing: B.Z., F.F, J.D., V-V. H., R.H., V.Z., J.M.H., J.A., M.S. and F.K.M.S

## Data availability

Representative tomograms containing the data shown in Figure 3, Figure S5 and Figure S9 have been deposited in the Electron Microscopy Data Bank under accession codes: EMD-18490, EMD-18491, EMD-18492, EMD-18493, and EMD-18494.

## Materials and Methods

### Cell culture, CDM growth, and CDM decellularization

Wild type *Homo sapiens* telomerase immortalized foreskin fibroblasts (TIFF) were cultured in Dulbecco’s modified Eagle’s medium (DMEM GlutaMAX, ThermoFisher Scientific, #31966047), supplemented with 20% (v/v) fetal bovine serum (ThermoFisher Scientific, #10270106), 1% (v/v) penicillin–streptomycin (ThermoFisher Scientific, #15070063), and 2% (v/v) 1 M HEPES (ThermoFisher Scientific, #15630080).

Cells were incubated at 37°C and 5% CO_2_ in a cell culture incubator. Phosphate buffered saline (PBS) used for all sterile cell culture work was purchased from ThermoFisher Scientific (#20012019).

For CDM growth, the protocol from (*31*) was adapted to use EM grids as substrate as follows: 150 or 200 mesh gold holey carbon grids (R 2/2, #N1-C16nAu15-01 and #N1-C16nAu20-01, respectively) were purchased from Quantifoil Micro Tools and glow discharged for 2 min in an ELMO glow discharge unit (Agar Scientific) prior to cell seeding.

Specimens were handled in a laminar flow hood from here on. EM grids were transferred to the lid of a sterile Falcon cell culture dish (Corning, #353004) with Parafilm stretched over its inside. All treatments described here were performed in this cell culture dish until otherwise stated. Grids were washed 1x with PBS and then coated with 20 µl of 50 µg/ml fibronectin (FN) (Sigma-Aldrich, #11051407001) in PBS for 1 hour at RT. Subsequently, grids were washed 2x with PBS and the FN was cross-linked for 30 min at RT with 20 µl of 1% (v/v) glutaraldehyde (Science Services, #E16220) diluted in PBS. After 3x PBS washes, any remaining glutaraldehyde was quenched with 20 µl of 1 M glycine (Lactan, #0079.2) in PBS for 20 min at RT. Grids were washed 1x in PBS and 2x in cell culture medium and then incubated for at least 15 min with cell culture medium prior to cell seeding.

TIFF cells were seeded onto grids in the cell culture dish in a drop of 20 µl with a density of 3.5 x 10^5^ cells/ml, resulting in ∼7,000 cells per grid. The seeded grids were incubated for 1-2 h in the cell culture incubator. During this incubation, 3D printed cubic grid holders (*36*) (see below) were washed 1x with PBS and 2x with cell culture medium and then incubated in cell culture medium for at least 1 h in a well in a 24-well plate (TPP, #92424). EM grids were then transferred into the grid holders and left there to incubate throughout CDM growth. Once the cells had grown into a confluent cell monolayer, typically within 2-3 days of seeding, the medium was exchanged every other day with new cell culture medium supplemented with 50 µg/ml ascorbic acid (Sigma-Aldrich, #A92902) and 10 mM HEPES.

Throughout all cell culture experiments, Dumont tweezers, medical grade, style 5 and style 7 were used.

### 3D printing of grid holders

Grid holders are described in detail in (*36*). Square base grid holders were generated using either a Green-TEC Pro filament (3D Jake) or PETG (Filament PM) with a printing resolution of 0.2 mm layer height for the first layer and 0.15 mm for all additional layers. All printing was done using a 0.4 mm nozzle. Stringing was removed from the grid holders after printing and all holders showing printing errors were discarded. Green-TEC Pro grid holders were washed 1x with perform® classic alcohol EP for 30 min and 2x with distilled H_2_O and then subsequently autoclaved prior to every use. PETG grid holders were treated identical, but were sterilized using 20 min of UV irradiation instead of autoclaving. Grid holders were re-used up to 15 times and stored under sterile conditions until use.

### CNA35-EGFP for CDM live-staining

The bacterial expression vector for pET28a-EGFP-CNA35 was a gift from Maarten Merkx (Addgene plasmid # 61603; http://n2t.net/addgene:61603; RRID:Addgene_61603) (*50*). BL21 *Escherichia coli* cells were used for expression and induced with 0.1 mM IPTG (Thermo Scientific, #R0393) at an OD600 of 0.6. The construct was expressed at 37°C for 4 h under constant agitation and cells were then pelleted by centrifugation with 6,000 g for 15 min at RT. The cell pellet was resuspended in a freshly prepared buffer containing 20 mM Tris (Lactan, #9090.3), 500 mM NaCl (Lactan, #P029.2), 5% (v/v) Glycerol (Sigma-Aldrich, #G5516-500ML), 2 µM ZnCl_2_ (Carl Roth, #3533.1), 1 mM PMSF (Sigma-Aldrich, #P7626-1G), 1 mM TCEP (Lactan, #HN95.2), pH 8.0. The resuspended cell pellets were snap-frozen in LN_2_ and stored at −80°C until purification.

Cells lysis was achieved through three cycles of freeze/thaw for 20 min at −80°C and 42°C, respectively. Cell debris was removed by centrifugation at 50,000 g for 1 h at 4°C. Subsequently, 10% PEI (Polysciences, #24966-100) was added to a final concentration of 0.3% to precipitate nucleic acid. Additionally, ammonium sulfate (Millipore Sigma, #1012115000) was added to a final concentration of 40% to precipitate proteins while stirring overnight at 4°C. Precipitated proteins were pelleted by centrifugation at 6,000 g for 10 min at 4°C and then dissolved in 20 mM Tris, 500 mM NaCl, 2 mM TCEP, and 20 mM Imidazole (Sigma-Aldrich, #56750), pH 8.0, while stirring at 4°C for 30 min.

CNA35-EGFP was purified from this solution through the application to a nickel sepharose column, HisTrap FF 1 mL (Cytiva, #17531901). After application, the column was washed with washing buffer (20 mM Tris, 500 mM NaCl, 2 mM TCEP, 20 mM Imidazol, pH 8.0) and then elution buffer (20 mM Tris, 500 mM NaCl, 2 mM TCEP, 250 mM Imidazol, pH 8.0). Fractions containing protein were pooled and dialyzed at 4°C overnight against dialysis buffer (20 mM Tris, 500 mM NaCl, 0.5 mM TCEP, pH 8.0). Aliquots of the purified protein were flash-frozen in LN_2_ and stored at −80°C until use.

CNA-EGFP was diluted to a final concentration of 1 µM in cell culture medium for all live staining of CDMs. For staining, CDMs were washed 1x with the staining solution and then incubated with staining solution for 1-2 h in the cell culture incubator. After staining, CDMs were washed 3x with cell culture medium and kept in the cell culture incubator and used for imaging and/or HPF within the next 1-3 h.

### Antibodies and stainings

4% PFA was diluted in PBS from a 16% stock solution (Science Services, #E15710) and pre-warmed to 37°C. Specimens were recovered from the grid holders and placed on Parafilm stretched over the inside of a cell culture dish lid, after which 4% PFA was added for fixation for 20 min at RT. Samples were washed 3x with PBS prior to an incubation in permeabilization solution (0.1% Triton X-100 [Sigma-Aldrich, #T8787] and 3% BSA [Sigma-Aldrich, #10735078001] in PBS) for 5 min at RT. After washing 3x gently with PBS, specimens were incubated in blocking solution (3% BSA in PBS) for 1 h at RT. Subsequently, blocking solution was removed and the sample was incubated in anti-fibronectin I antibody from rabbit (Sigma-Aldrich, #F3648), diluted 1:500 in blocking solution, overnight at 4°C in a wet chamber.

The next morning, specimens were washed 3x with PBS before incubation in a solution of anti-Rabbit-IgG-ATTO 594 (Merck, #77671-1ML-F) as secondary antibody, phalloidin-ATTO 488 (ATTO-TEC, #AD488-81), and 4’,6-Diamidino-2-phenylindole dihydrochloride (DAPI, Sigma-Aldrich, #32670-5MG-F), all diluted 1:500 in blocking solution. Samples were incubated for 1-2 h at RT in a wet chamber in the dark. After this incubation, samples were washed 3x with PBS and stored at 4°C in a wet chamber in the dark until imaging.

### Light Microscopy

Specimens stained with EGFP-CNA35 were kept on Parafilm in a drop of cell culture medium and quickly assessed for sample integrity and quality prior to vitrification. Fixed specimens stained with antibodies and other dyes were kept on Parafilm in a drop of PBS. For this assessment we used a Zeiss LSM800 microscope with a Plan-Apochromat 20x / NA 1.0 W DIC water-dipping (WD = 1.8mm) objective. Z-stacks with 1 µm steps over the whole height of the specimen, from the EM substrate to the top of the CDM, were acquired using the ZEN 2.6 software. Typically, at least 3 positions per specimen were acquired. Live samples were kept at 37°C for the duration of the imaging process and returned to a cell culture incubator after a maximum of 30 min. Alternatively, a Zeiss Axioscope with a W N-Achroplan 20x/0.5 water-dipping (WD = 2.6 mm) objective was used for quick assessments prior to HPF. A maximum intensity Z-projection was applied to Z-stacks using Fiji (*51*). To improve the visibility of these images, contrast and brightness were adjusted as necessary.

During this initial assessment, any specimens showing distortions of ECM fibers, damage to the CDM, or bending of the EM grids were removed from the sample pool.

### Mass Spectrometry

#### Sample processing

TIFF cells were seeded on 10 cm diameter cell culture dishes (Sarstedt, #83.3902) and treated as described above for CDM growth. Cell cultures dishes were not coated with FN prior to cell seeding to prevent the introduction of a bias to the mass spectrometry analysis. CDMs were grown for 14 days with ascorbic acid treatment every other day.

CDMs were then decellularized following the protocol published by Kaukonen *et al.* (*31*). Briefly, CDMs were washed with extraction buffer consisting of 0.5% Triton-X100 and 0.56-0.6% Na_4_OH (Sigma-Aldrich, #221228-1L-A) in PBS that had been prewarmed to 37°C. The washes were repeated until the fibroblasts were extracted, as observed by phase contrast microscopy on a standard cell culture stereo microscope. Typically, this process takes up to 5 min and 3-4 washes for TIFF CDMs. After cells were extracted, CDMs were washed with PBS and treated for 1 h at 37°C with a buffer of 50 µg/ml DNaseI (Roche, #11284932001), 5 mM MgCl_2,_ and 1 mM CaCl_2_ diluted in PBS. Specimens were then washed three times with PBS and immediately fixed with 4% PFA in PBS. After decellularization, CDMs were scraped off the cell culture dish surface with a cell scraper (Sarstedt, #83.1830) and the matrix was transferred into 1.5 ml centrifugation tubes.

Subsequently, CDMs were centrifuged (13,000 g, 5 min), supernatants were removed, and the resulting pellets were resuspended in a buffer consisting of 100 µl 8 M Urea (Sigma-Aldrich, #U1250), 100 mM TEAB (triethylammonium bicarbonate; Sigma-Aldrich, #T7408-100ML), and 25 mM TCEP (tris(2-carboxyethyl)phosphine hydrochloride, ThermoFisher Scientific, #77720). Samples were then sonicated (Bioruptor plus, Diagenode, 10 x 30s/30s ON/OFF cycles) and heated up to 37°C for 2 h while shaking (800 rpm, Thermomixer F1.5, Eppendorf). Following this, specimens were alkylated by the addition of 100 µl 50 mM Iodoacetamide (ThermoFisher Scientific, #A39271) and incubated for 30 min in the dark while shaking (800 rpm). Then, 200 µl 100 mM TEAB was added to the specimens and they were digested with 10 µl PNGase (Gibco, #A39245) at 50°C for 2 h.

Adapting the protocol published in (*52*), the sample was diluted by addition of 390 µl 100 mM TEAB, then supplemented with 8 µL Trypsin/LysC (1 µg/µl; Promega, #V5072) and digested overnight at 37°C. Following this, 4 µl Trypsin/LysC (1 µg/µl) was added to the sample and it was incubated for a further 2 h, then 90 µl 10% TFA (trifluoro-acetic acid; ThermoFisher Scientific, #10723857) was added for acidification. A tC18 SepPak plate (Waters, #1860002318) was used for clean-up according to the manufacturer’s protocol.

#### LC-MS/MS analysis

The sample was dried, re-dissolved in 0.1 % TFA, and analyzed by LC-MS/MS on an Ultimate High Performance Liquid Chromatography (nano HPLC, ThermoFisher Scientific) coupled to a Q-Exactive HF (ThermoFisher Scientific). An Acclaim PepMap C18 trap-column (5 µm particle size, 0.3 mm ID x 5 mm length; ThermoFisher Scientific, #160454) was used to concentrate the sample, which was then bound to a 200 cm C18 µPAC column (micro-Pillar Array Column; PharmaFluidics, #5525031518210B) and finally eluted with a constant flow of 600 nl/min over the following gradient: solvent A, 0.1% formic acid (ThermoFisher Scientific, #160454) in water; solvent B, 80% acetonitrile (ThermoFisher Scientific, #10001334) and 0.08% formic acid in water. Step 1: 5 min of 2% solvent B. Step 2: 160 min of 31% solvent B. Step 3: 185 min 44% solvent B. Mass spectra were acquired in positive mode with a Data Dependent Acquisition method: Full Width at Half Maximum (FWHM) 120 s, Mass Spec scan acquired without fragmentation parameters (MS1): Profile, 1 microscan, 120,000 resolution, Automatic Gain Control (AGC) target 3e6, 50 ms maximum IT, 380 to 1500 m/z; up to 20 MS2s per cycle. Mass Spec scan acquired after one round of fragmentation (MS2) parameters: Centroid mode, 1 microscan, 15,000 resolution, AGC target 1e5, 20 ms maximum IT, 1.4 m/z isolation window (no offset), 380 to 1500 m/z, NCE 28, excluding charges 1+, 8+ and higher or unassigned, 60 s dynamic exclusion.

#### Data analysis

Raw files were searched in MaxQuant 1.6.17.0 against a *Homo sapiens* reference proteome downloaded from UniProtKB. Fixed cysteine modification was set to Carbamidomethyl. Variable modifications were Oxidation (M), Acetyl (Protein N-term), Deamidation (NQ), Gln->pyro-Glu, Phospho (STY), and Hydroxyproline. Match between runs, dependent peptides, and second peptides were active. All False Discovery Rates (FDRs) were set to 1%. MaxQuant results were further processed in R using in-house scripts, which, starting from MaxQuant’s evidence.txt (PSM) table, perform parsimonious protein groups inference and generate an Excel-formatted protein groups table. GO annotations were downloaded from UniProtKB. ECM proteins were defined as proteins annotated with GO term “Extracellular Matrix” (GO:0031012) or any of its descendants.

ECM proteins were then sorted according to the normalized log10 of the estimated protein group expression value and listed in **Table S2**.

### Array Tomography

Sample preparation was done according to the OTO fixation protocol described in Deerinck *et al.* (2010) in order to enhance the contrast of sample for SEM (*53*).

TIFF CDMs were grown on glass cover slips (Epredia, #CB00150RA120MNZ0) for 14 days with ascorbic acid treatment as described in the section “**Cell culture, CDM growth, and CDM decellularization”** and then fixed with 2% PFA and 2.5% glutaraldehyde in 0.1M PB for 1 h at RT. Samples were washed with 0.1 M PB and contrast was enhanced using 2% aqueous osmium tetroxide (Science Services, #E19110) and 1.5% potassium ferrocyanide (Sigma, #P9387-100G) in 0.1 M PB for 30 min in the dark. After washing with MilliQ water, the samples were incubated in thiocarbohydrazide (Sigma, #88535-5G) for 20 min at RT and subsequently washed with MilliQ water.

Following this, specimens were placed in 2% aqueous osmium tetroxide for 30 min at RT in the dark and then washed again with MilliQ water. Samples were then incubated overnight in 1% aqueous uranyl acetate (AL-Labortechnik, #77870.02) at 4°C. The following morning, they were again washed with MilliQ water, incubated in Walton’s lead aspartate solution prepared from L-aspartic acid (Sigma, #A8949-25G) and lead nitrate (Sigma, #228621-100G) for 30 min at 60°C, and washed again in MilliQ water.

Samples were subjected to a graded series of ethanol of 50%, 70%, 90%, and 2x 100% (Bartelt, #32221-2.5L) for dehydration and then placed in anhydrous acetone (Bartelt, #CL0001722500). They were then infiltrated in a graded series of hard DurcupanTM ACM resin (Sigma, Component A: #44611-100ML, B: #44612-100ML, C: #44613-100ML, D: #44614-100ML) in acetone, and placed in pure Durcupan overnight. The following day, the coverslips were put on an ACLAR® foil (Science Services, #E50425-10), and a BEEM® capsule (size 00, Science Services, #E70020-B), filled with fresh resin, was placed upside down in the middle of each coverslip. Samples were placed in a 60°C oven for 3 days for polymerization. After this, they were dipped in LN_2_ until the coverslip could be carefully removed with a razor blade.

Samples were trimmed with an Ultratrim diamond knife (Diatome) using an Ultramicrotome EM UC7 (Leica Microsystems). A carbon-coated 8 mm wide Kapton tape (RMC Boeckeler) was plasma treated using an ELMO glow discharge unit, equipped with a homemade reel-to-reel motorized winder, to increase its hydrophilicity. Serial ultrathin sections of 70 nm thickness were cut with a 4 mm Ultra 35 diamond knife (Diatome) and lifted up with the plasma-treated tape using an automated tape-collecting Ultramicrotome ATUMtome (RMC Boeckeler).

The tape used to collect the serial sections was cut into strips and mounted on a 4-inch silicon wafer (University Wafer) with conductive double sided adhesive carbon tape (Science Services, #P77819-25). The wafer was then coated with a 5 nm carbon layer using an EM ACE 600 (Leica Microsystems) to ensure conductivity. The collected sections were then imaged on a Field Emission-SEM Merlin compact VP (Carl Zeiss) equipped with the Atlas 5 Array Tomography software. The high-resolution serial images for 3D-SEM reconstruction were taken with 10 nm pixel resolution at 5kV using a backscattered electron detector.

Serial SEM images were down-sampled to approximately isotropic resolution (x,y: 80 nm, thickness 70 nm). These images were then aligned using a custom Matlab script: The optimal affine transformation linking consecutive images was found employing an evolutionary optimizer based on pairwise SURF features, employing the M-estimator sample consensus algorithm and using mean squares as a quality metric.

The process of pixel classification for cell bodies, nuclei, and ECM was executed using the auto-context workflow in Ilastik (version 1.4.0) (*54*). The classification of filamentous structures was done separately via the pixel classification workflow in Ilastik. The output generated was subsequently imported into Imaris (version 9.3) for visualization and reconstruction of cell body and nuclear surfaces.

To visualize the alignment of the cell/nucleus major axis with the fiber orientation the Fiji plugin OrientationJ was used (settings: σ=16, gradient= Cubic Spline) (*55*).

### High pressure freezing (HPF)

Carriers were designed to fit the Z-height of on-grid CDMs and custom-produced at the ISTA Miba Machine Shop. Two types of 3 mm diameter carriers were combined for HPF of CDMs: Carriers of type A had a height of 0.5 mm, with a 2 mm diameter recess of a depth of ∼20 µm (±5 µm machining inaccuracy). Carriers of Type B had height of 0.5 mm without any recess. To ensure proper carrier sandwich height after assembly, every single carrier was measured manually with a digital Vernier caliper for its height, and any carrier with more than ±20 µm derivation in height was removed.

All carriers were cleaned by three rounds of sonication in pure ethanol and then dried at 60°C on a hot plate. Prior to use, carriers were fully coated with 1-hexadecene (Sigma, #H2131-100ML). CDMs were incubated in the used cryoprotectants listed in **Table S3** 30 min prior to vitrification and kept at 37°C, 5% CO_2_ during this incubation time. CDMs were kept at 37°C up until HPF carrier sandwich assembly and then frozen as quickly as possible with a BAL-TEC HPM010. To avoid excess contamination, specimens were stored in cryo-vials in LN_2_ directly after HPF and then transferred to a freshly cooled, clean clipping station for disassembly. Recovered specimens were clipped into FIBSEM AutoGrids (ThermoFisher) marked as described in Wagner *et al.*, 2020 (*56*) and stored until further use.

The cryoprotectants listed in **Table S3**, Dextran (#31389-100G), Sucrose (#84100-1KG), Polyvinylpyrrolidone (PVP, #PVP10-100G), and BSA (#10735078001), were purchased from Sigma-Aldrich.

### Cryo-fluorescence microscopy

All specimens were screened on a Leica EM Cryo CLEM microscope (Leica Microsystems) using the Leica Application Suite 3.7.0. The LasX navigator was used to acquire tile scans of entire FIBSEM Autogrids to facilitate correlation for FIB milling. Z-stacks of regions of interest were acquired to assess presence of collagen fibers and to select the best positions for ion-beam milling. All specimens showing damage to the CDM or strong distortions of the EM grid were discarded after imaging.

### Cryo-FIB milling

Cryo-lift out lamellae were generated using a second generation Aquilos (Aquilos II) instrument (Thermo Scientific). The instrument was operated using the xT user interface and the MAPS 3.14 software (TFS). The FIB was operated at 30kV, and the milling progress was monitored using the SEM beam at 25 pA and 2-5 kV.

After sample loading, overview maps of the high-pressure frozen specimens were acquired and correlated with the images obtained on the Leica EM Cryo CLEM microscope to identify regions of interest. The milling slot on the FIBSEM AutoGrid was used for improving correlation by using its rim, visible in both reflected light microscopy and SEM, as landmark.

Then specimens were sputter coated with platinum for 30 s at 30 mA and 10 Pa with the built-in sputter coater, followed by a GIS deposition of 1.5-2 µm metalorganic platinum. Another tile-scan overview image was then taken using the MAPS software.

Lift-out sites were identified by CLEM and set to eucentric position. Steps performed for the lift-out FIB milling will be explained using **Figure S2C** as illustration. **Figure S2C-1**: Trenches for the lift-out procedure were cut at a stage tilt of 7° and a relative stage rotation of 180° to position the FIB perpendicular to the sample. The trenches in front, behind, and to the side of the lift-out were milled in cross-section (CS) patterns with 3 nA and their size was adjusted as needed. The front and back of the lift-out were polished smooth by milling with cross-section-cleaning (CSC) patterns fitted to the width of the lift-out. **Figure S2C-2**: Undercuts were performed at 28° stage tilt, at the default stage rotation, with 1 nA. Rectangle milling patterns were placed below and to either side of the lift-out, leaving it attached to the bulk sample only on a small anchor, marked red. The micromanipulator needle was then attached by redeposition milling, using a CS pattern below and above the needle set to 0.5 µm z-depth, with a Multi-Pass setting of 1 at 0.5 nA. Pattern placement is shown in blue in the Figure. After visual confirmation of successful attachment, the remaining anchor to the bulk sample was removed at 0.5 nA with a rectangle pattern placed at the anchor position, as indicated by the red pattern in the Figure. **Figure S2C-3 and S2C-4**: The lift-out was then lifted up and out of the bulk sample and transferred to the second shuttle position, which held a half-moon grid (Ted Pella, #10GC04), with 4 finger-like extensions for lift-out attachment. **Figure S2C-5**: The lift-out was attached to a finger by redeposition milling, using four CS patterns at 0.5 nA with a Multi-Pass setting of 1, a z-depth of 3 µm, and a surface of ∼2×2 µm. The pattern placement is indicated in blue in the Figure. Following visual confirmation of the attachment, the needle was pulled off gently to the side, so it could be directly reused for the next lift-out without any necessary cleaning steps. **Figure S2C-6**: The portion of the lift-out that was used for the needle-attachment was then removed by placing a rectangle milling pattern and FIB milling with 1 nA, as indicated by a red pattern in the Figure. **Figure S2C-7 to S2C-9**: Each lift-out was then milled down with rectangle patterns to ∼200-250 nm thickness in several steps, reducing lamella width and milling current in each step, resulting in a symmetric stair-like anchor. Lamellae were thinned down to 3 µm thickness with 1 nA, then to 1.5 µm thickness with 0.5 nA, and to 900 nm using 0.1 nA. Here, the stage was tilted to +/− 1° and the lamella overhangs above and below were milled with 0.1 nA to facilitate a parallel shape of the lamella rather than a wedge-like one. The lamella was then milled down to 500 nm with 50 pA and then again lamella overhangs were removed with +/− 0.5° tilts. In a final step, the lamella was thinned down to 200 nm with a milling current of 30 pA.

All samples were stored in autogrid boxes in liquid nitrogen until TEM imaging.

### Cryo-TEM and cryo-ET

A TFS Titan Krios G3i operated at 300 kV in nanoProbe energy-filtered transmission electron microscopy (EFTEM) mode equipped with a Gatan K3 BioQuantum direct electron detector with a slit width of 20 eV was used for data acquisition on cryo-lift out lamellae. Zero Loss Peak (ZLP) and energy filter tuning were done using DigitalMicrograph 3.42 (Gatan). Coma-free alignment and astigmatism correction were performed using SerialEM 4.0 beta5 (*57*). For medium magnification images, a pixel size of 13.74 Å at a nominal magnification of 6,500x was used.

For tilt series acquisition, the camera was operated in counting mode using hardware binning and dose fractionation, with 8 frames per tilt. The total dose applied was 180 e/Å^2^, divided into 61 images (for a 2° increment tilt scheme). All tilt series were acquired with a dose-symmetric scheme starting from the lamella pre-tilt angle in a range of −60° and +60° (*58*) at a defocus of −8 µm. The pixel size was set to 2.137 Å at a nominal magnification of 42,000x. Datasets were acquired using SerialEM (*59*) and PACE-tomo (*60*).

Tomograms were reconstructed using weighted back-projection and patch tracking, employing the IMOD software (*61*) as well as AreTomo (*62*), with a binning of 8. Bad tilts compromised by movement, beam obstruction, or strong reflections were removed. A SIRT-like filter (equivalent of 15 iterations) was applied during tomogram reconstruction for selected tomograms to visualize the collagen banding pattern more clearly. Tomograms were additionally processed with IsoNet (*63*) to increase the signal-to-noise ratio (SNR) and fill in the missing-wedge information. For this, raw bin 8 tomograms were deconvoluted with IsoNet and subsequently used to train the neural network for 50 iterations in batches of 10 tomograms. The mask patch size used for training was set to 6 subtomograms with a cube size of 64 pixels. The same tomograms were then also used for the reconstruction of the missing-wedge information and improvement of the SNR. All data from tomograms shown in this paper originate from these IsoNet filtered tomograms unless otherwise stated.

### Segmentation

Tomograms were segmented in the Dragonfly software, Version 2020.2 for Linux (Object Research Systems (ORS) Inc, Montreal, Canada, 2020). For each tomogram, four slices were selected in the Segmentation Wizard and manually segmented. These segmented slices were then used for the training of a deep learning model utilizing a U-Net architecture, with an input dimension of 2.5D (number of slices = 3), a depth level of 5 and an initial filter count of 128. Following this training, the generated model was used to segment the whole tomogram and corrections were manually applied as necessary. Each class of object was extracted as a ROI from the segmentation and rendered into a 3D contour mesh. Following this, the mesh was smoothened using the Laplacian Smoothing method with 2-5 iterations as needed, and the smoothened contour mesh was exported as a .stl file for visualization in UCSF ChimeraX (*64*).

The volume shown in Figure 3A was partially segmented in the Amira-Avizo software, version 2020.2 (Thermo Fisher Scientific): Plasma membranes and vesicles were tracked manually in Amira. All manually segmented structures were exported from Amira as .mrc files and then imported into ChimeraX, smoothened, and visualized together with the files imported from Dragonfly. All objects were rendered for display using ChimeraX.

To display all objects on the same scale in ChimeraX, Dragonfly files were first upscaled to match the tomogram dimensions. Where there were obvious defects from Dragonfly export, model meshes were manually fixed using Blender 3.5.1. All models were loaded into ChimeraX and positions of .stl objects manually positioned relative to the tomogram. Amorphous density in the extracellular matrix was generated by first inverting density for each tomogram then using the “Segger” tool in Chimera 1.17.1 (*65*). Regions of density inside of cells were manually excluded, and the remaining density in the ECM was extracted from the volume for rendering in ChimeraX.

Final figures and movies were generated using ChimeraX. Camera perspective was set to mono, lighting depthCueStart and depthCueEnd were both set to 1. Lighting and silhouettes were otherwise set to default settings using the “soft” lighting preset. A volume box outline was colored as “grey” and displayed for the amorphous matrix only. The amorphous matrix was thresholded manually to yield a result representative of the raw tomogram, and Gaussian filtered to a value of “22” within Chimera 1.17.1. Amorphous matrix was colored #FFDAB9 (peach puff) with transparency set to 40% for images and 0% for movies. Plasma membranes were colored #6495ED (cornflower blue), vesicles colored #87CEEB (sky blue), collagen colored #3CB371 (medium sea green), small ECM filaments colored #FFFF00 (yellow), microtubules colored #FF8C00 (dark orange). Actin was colored #00F0F0 (cyan), intermediate filament colored #DC5A9B (magenta), and granules colored #8FB38D (light green). Tomogram slices were overlaid with the segmentation using ArtiaX version 0.3 plugin (*66*). Slice representation was set to “plane” in volume viewer. Tomograms were manually brightened within ArtiaX for display purpose.

## Supplementary Information

**Figure S1:**
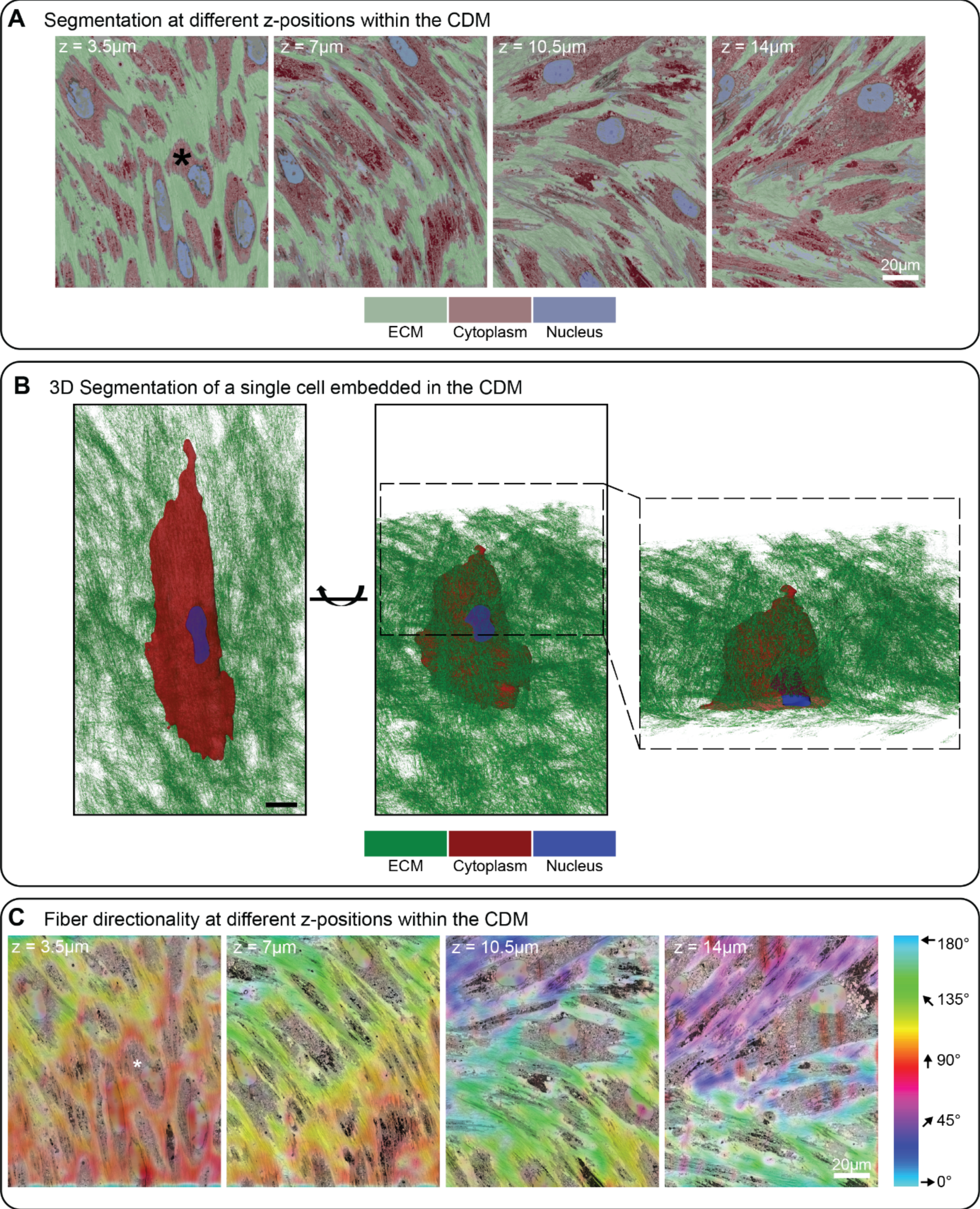
Room-temperature SEM array tomography of TIFF CDMs grown for 14 days. **A)** Four exemplary 70 nm thick sections from different z-height positions within the CDM are shown. Nuclei, cytoplasm, and ECM were segmented using Ilastik (*54*) and are colored as indicated. **B)** 3D segmentation of an exemplary cell embedded within the CDM (denoted by a black asterisk in A), shown in a top (left) and oblique view (middle). On the right side, an ortho-slice view of the same cell is shown, highlighting the thin z-height dimension of the cell embedded within the ECM. Please note that the other cells in vicinity of the segmented cell have been omitted from this view to facilitate visualization. Color code of the segmentation as in A). **C)** The directionality of cells and fibers is color-coded as indicated on the right, corresponding to the angles of cells and fibers. The same z-positions as shown in A) are depicted to show the change in directionality throughout the height of the CDM. Scale bar dimensions are annotated in the figure.

**Figure S2:**
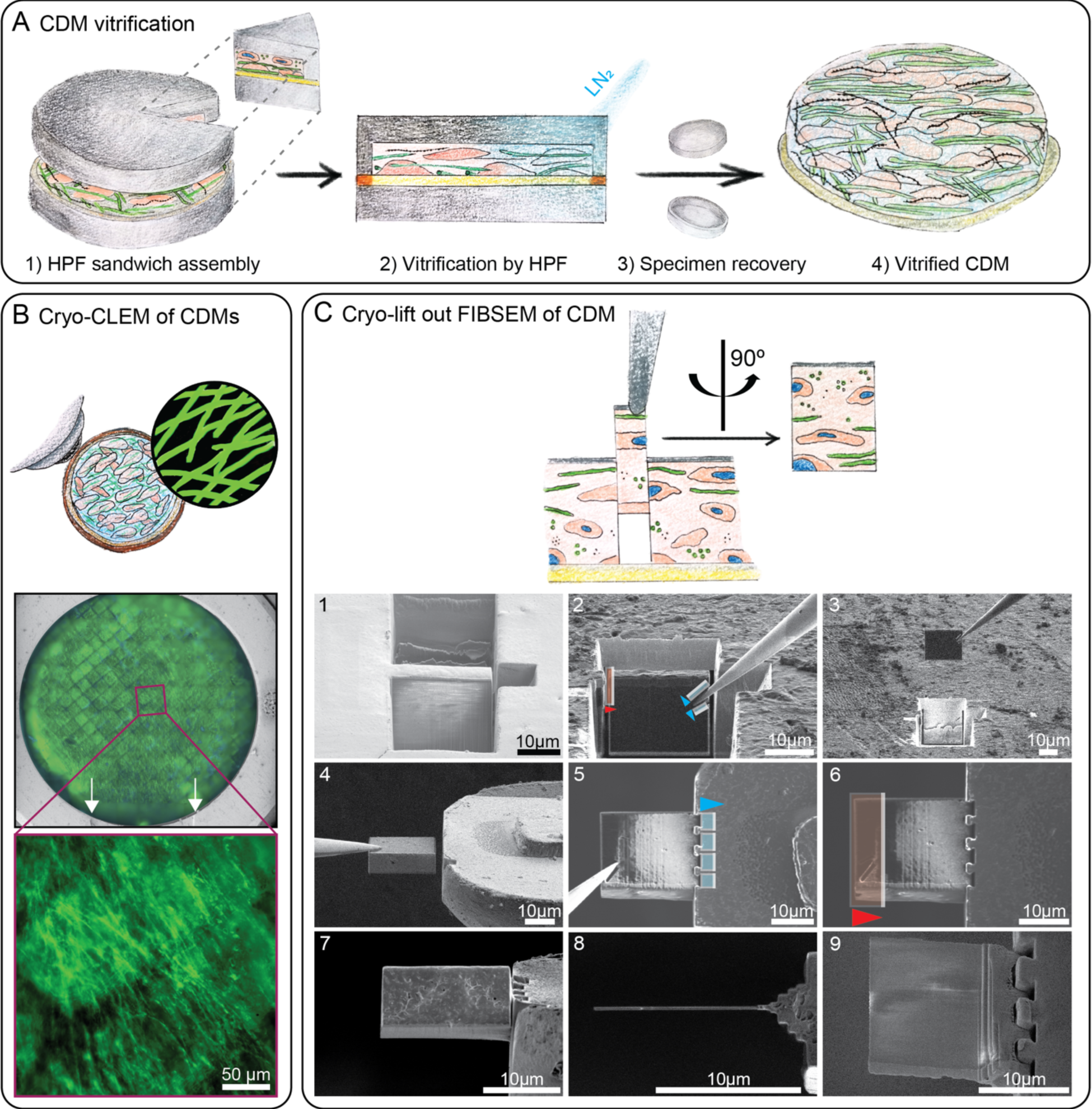
HPF and cryo-lamella preparation of CDMs for cryo-ET. **A)** Schematic workflow of on-grid CDM vitrification: 1) sandwich assembly of on-grid CDMs, live-stained for collagen; (2) vitrification via HPF; (3) specimen recovery of vitrified CDM (4). **B)** All specimens are screened for their quality and to define regions of interest (ROI, annotated with a purple rectangle) by cryo-fluorescence light microscopy. A magnified image of the ROI, showing collagen fibers is depicted below. Reflected light is used to define landmarks for correlation, such as the milling window on the FIBSEM Autogrid, indicated by white arrows. **C)** Cryo-liftout FIBSEM workflow. Trenches are milled to isolate the lift-out (1), which is attached to a micromanipulator needle by redeposition milling (blue patterns, 2) and prior to cutting it loose from the bulk sample (red pattern, 2). The lift-out is extracted from the bulk sample (3) and attached to a finger-like protrusion on a half-moon grid by redeposition milling (4-5, blue patterns). The needle is cleanly pulled off and its attachment site is removed from the lift-out by FIB milling (red pattern) (6). The lift-out can be milled into a thin-lamella (7–9) compatible with cryo-ET. Arrowheads indicate milling direction. Scale bar dimensions are annotated in the figure.

**Figure S3:**
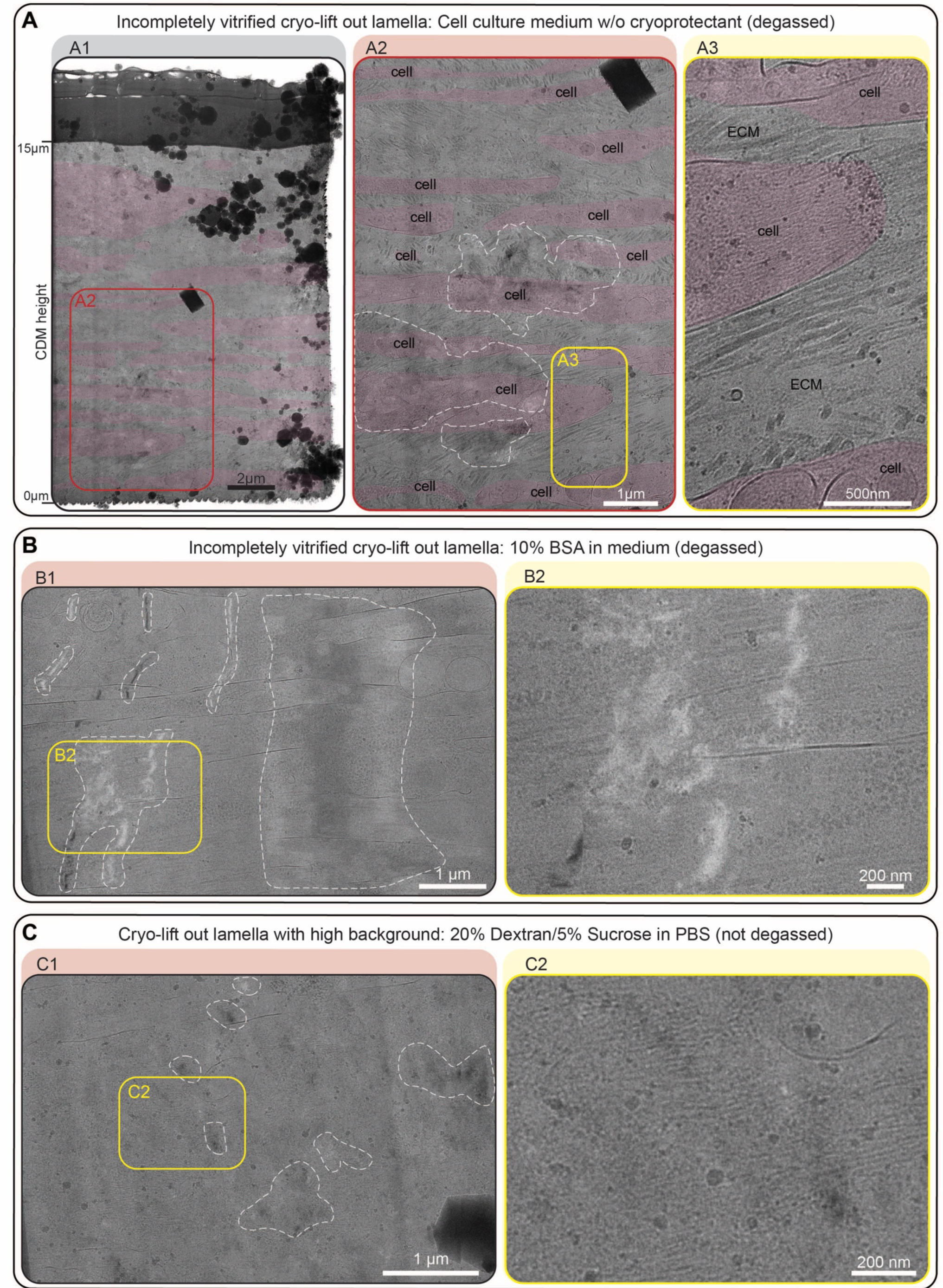
Examples of incompletely vitrified CDM lift-out lamellae. **A)** CDMs were vitrified in cell culture medium without cryoprotectants. **A1)** Overview of a whole cryo-lift out lamella. The lamella covers roughly 15 µm of CDM depth, ranging from proximal to the EM grid substrate (z = 0 µm) to close to the CDM surface (z = 15 µm). Cell bodies are annotated in transparent red color. **A2)** Zoom-in of the lamella as annotated with a red rectangle in A1. Areas with reflections caused by incomplete vitrification are marked with a white, dashed line. **A3)** Zoom-in of the area annotated with a yellow rectangle in A2, showing high contrast ECM structures of interest. **B)** Example of a lamella with incomplete vitrification, despite use of cryoprotectant (cell culture medium, with 10% BSA after the buffer has been degassed). **B1)** Lamella overview. Strong reflections are observed throughout the area (marked with white dashed line). **B2)** Zoom-in into the lamella, as annotated by a yellow rectangle in **B1**. The reflections obscure cell and ECM details, while the background would have been acceptable. **C)** Example of a lamella with too high background, introduced by the cryoprotection buffer (20% Dextran, 5% Sucrose in PBS, without degassing). **C1)** Lamella overview. Weak reflections can be seen throughout the area (white dashed line), resulting in a categorization of the vitrification status as incomplete. **C2)** Zoom-in into the lamella, annotated by a yellow rectangle in C1. The high background reduces visibility of cellular and ECM structures. Scale bar dimensions are annotated in the figure.

**Figure S4:**
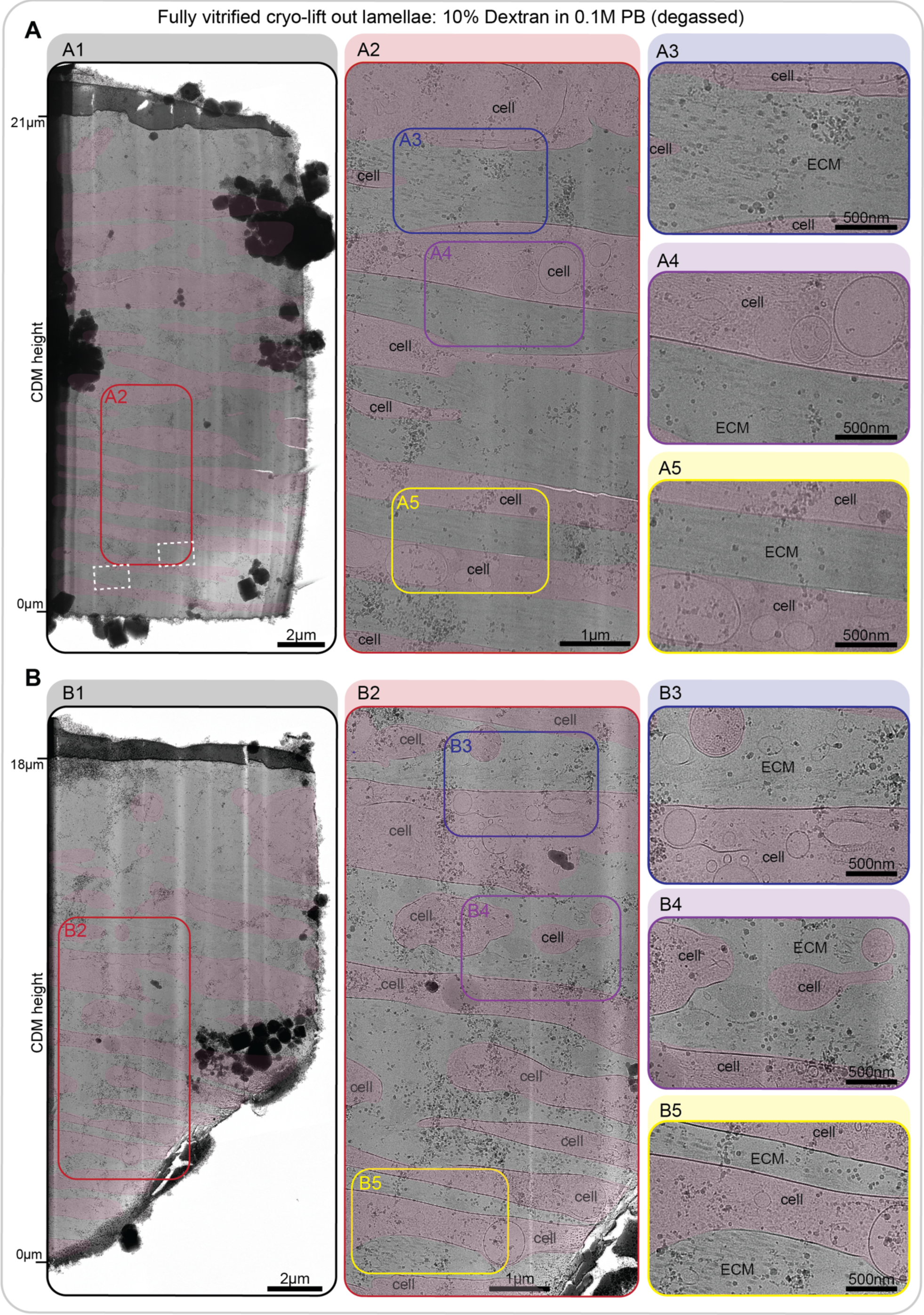
Additional examples of completely vitrified cryo-lift out lamellae. **A, B)** Two additional, completely vitrified lamellae of day 16 CDMs, high-pressure frozen in degassed 0.1M PB with 10% Dextran. White dashed rectangles denote the areas of acquisition for the tomograms shown in Figure S5. An overview map of each whole lamella is shown in **A1** and **B1**. The CDM height is indicated on the left, starting from 0 µm close to the EM grid and up to 21 and 18 µm at the surface of the CDM, respectively. Areas featuring structures of interest are depicted at higher magnification in **A2** and **B2.** Three ROIs are highlighted in each example and shown at higher magnification on the right (**A3-A5, B3-B5**). Cell areas are annotated with transparent red color, ECM areas are labeled as such. Scale bar dimensions are annotated in the figure. The experimental conditions of the shown CDMs are identical to the data shown in Figure 2.

**Figure S5:**
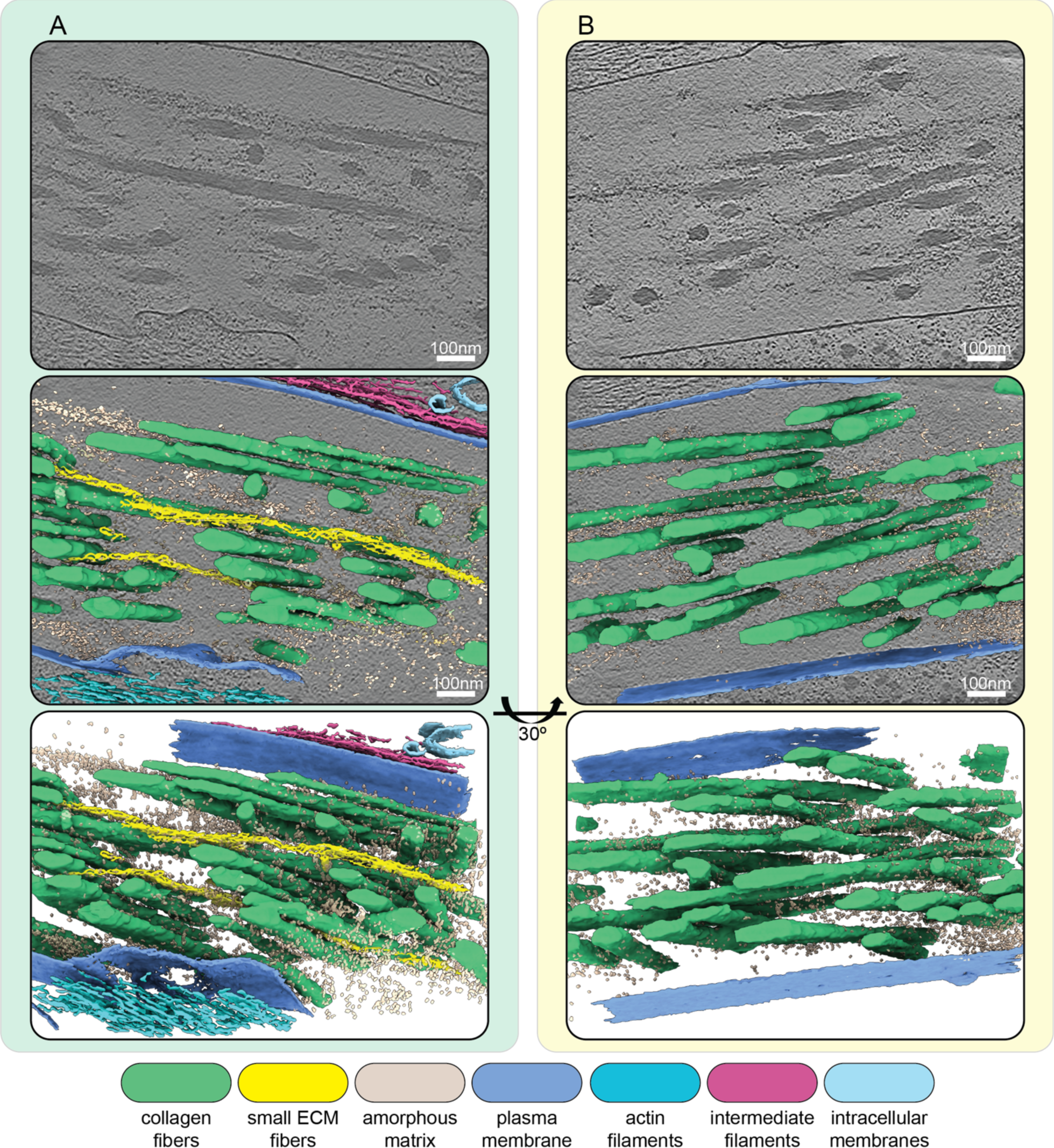
Additional examples of segmented cryo-electron tomograms. **A, B)** Segmentations of two exemplary IsoNet-processed tomograms acquired on the cryo-lift out lamella shown in Figure S4A. The top panels show a single central slice (thickness of 1.71 nm) of each respective tomogram. Cell and ECM fibers run in an oblique angle to the lamella, resulting in a side view of intra- and extracellular filaments. Middle panels show segmentations of tomograms overlaid over the tomogram slice. Bottom panels show an oblique view of just the segmentation volume. Scale bars indicate 100 nm. The color scheme for the different segmented cellular and ECM components is described in the figure.

**Figure S6:**
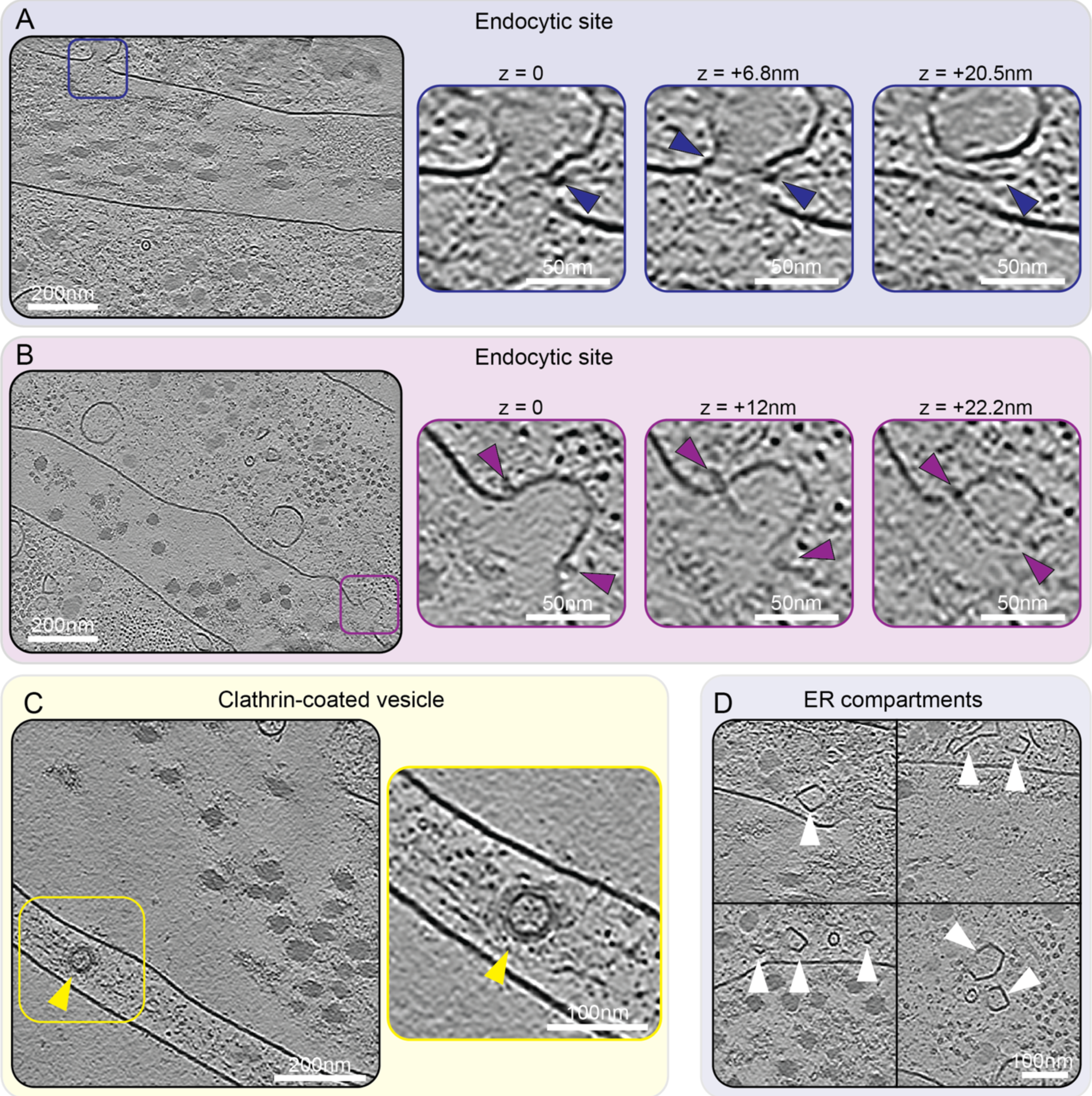
Membrane compartments – Endocytic sites, clathrin-coated vesicles, ER compartments. **A, B)** Single slices of two IsoNet-processed tomograms are shown on the left. Both tomograms show an endocytic site, highlighted by blue (A) and purple (B) squares, respectively. For both tomograms, these sites are shown at higher magnification and at different z-positions to better illustrate the structural details. Enodcytosis-associated proteins, potentially BAR-domain proteins, can be seen assembled at the neck of the forming vesicle, indicated by blue and purple arrowheads. The difference in z-position is indicated above each image. **C)** A single central slice of an IsoNet-processed tomogram shows a clathrin-coated vesicle annotated by a yellow arrow head. **D)** Four exemplary regions containing ER compartments (annotated by white arrowheads) in IsoNet-processed tomograms are shown. All tomogram slices have a thickness of 1.71 nm. Scale bar dimensions are annotated in the figure.

**Figure S7:**
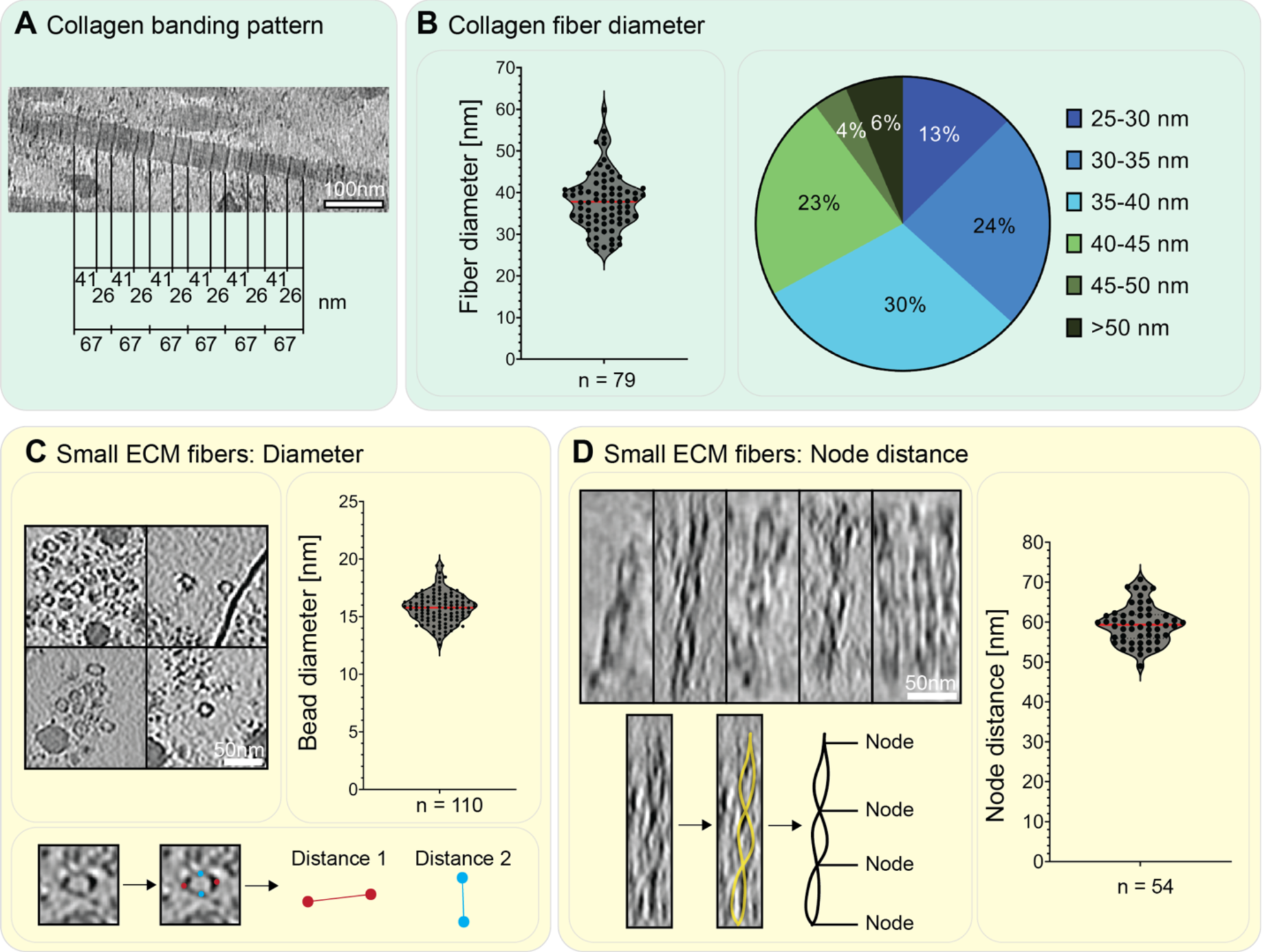
Details of ECM fibers. **A)** Focused view on the banding pattern of collagen fibers. The shown tomogram was reconstructed using a SIRT-like filter to enhance the visibility of the banding pattern. The typical collagen repeat pattern of a 41 nm wide band followed by a 26 nm narrow band, summing up to an overall repeat of 67 nm, is indicated. **B)** Fiber diameter of 79 collagen fibers as measured in cross-section view (to reduce missing-wedge effect-caused inaccuracies). Each single measurement is shown as a dot in a violin plot. A red line indicates the median. On average, collagen fibers have a diameter of ∼38 nm (SD = 6.5 nm), ranging from 26 to 60 nm. The distribution in diameter of the measured fibers is shown in more detail in the pie-chart on the right. **C-D)** Characterization of small ECM fibers by measuring their diameter (C) and node-to-node repeat distance (D) in IsoNet-corrected tomograms. **C)** Four exemplary images showing one or more small ECM fibers (left). Their diameter has been measured at their widest point. As fibers were not round, the average of two distances, measured perpendicular to each other, was calculated. 110 individual measurements are displayed in the respective violin plot shown on the right, with the red line indicating the median. The average bead diameter is 15.74 nm (SD = +/− 1.31). **D)** The node-to-node distance of the small ECM fibers was determined for 54 individual measurements. Exemplary images of IsoNet-corrected tomogram regions used for measurements are shown on the left. Below, one exemplary filament is traced in yellow, with the tracing shown as standalone in black next to it. The measurements are summarized in the violin plot on the right, the red line indicates the median. Overall, the node distance is at average 60.30 nm (SD = +/− 7.82 nm). Scale bar dimensions are annotated within the figure.

**Figure S8:**
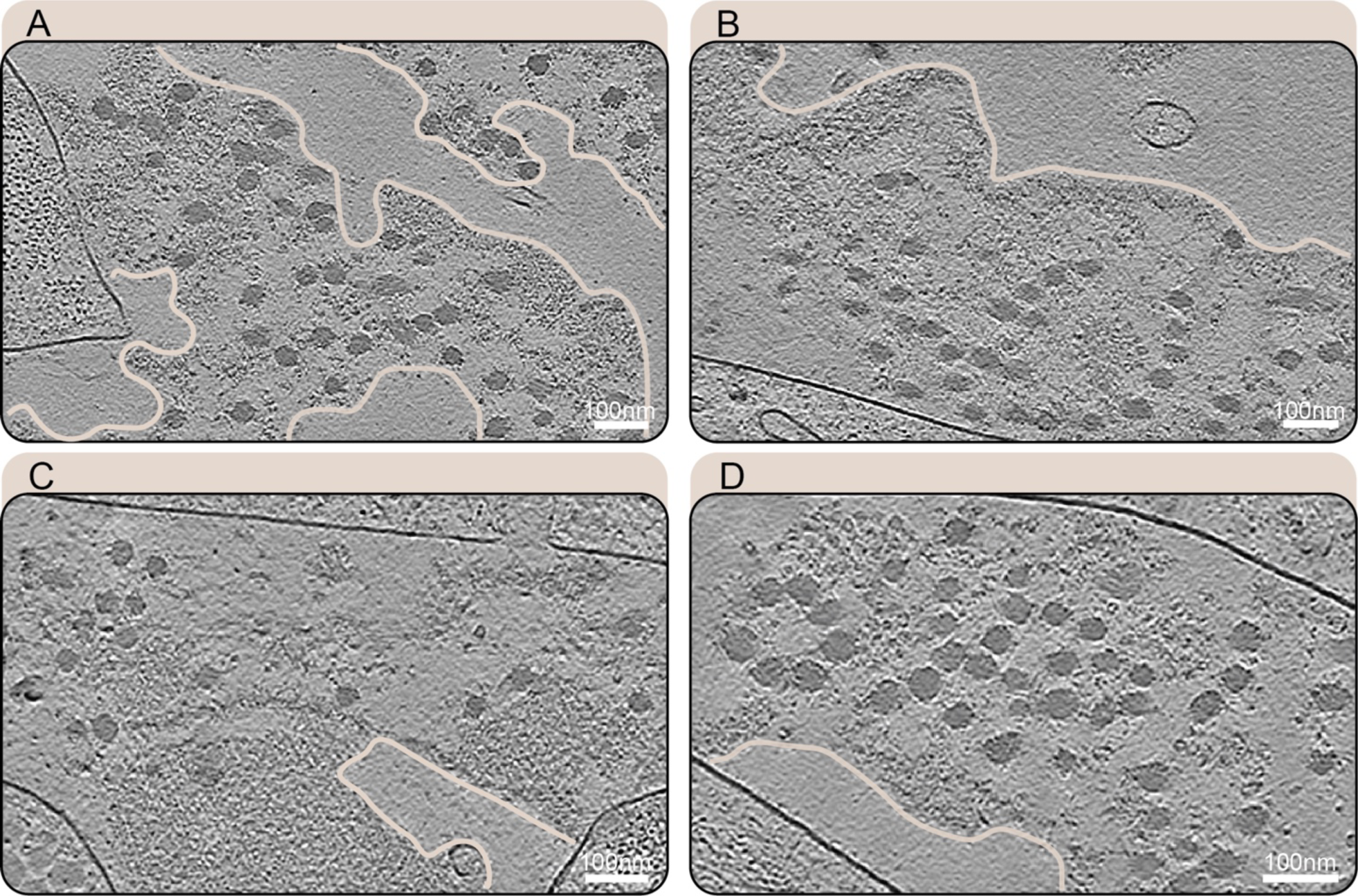
Amorphous matrix details. **A-D**) Slices (1.71 nm thickness) through IsoNet-processed tomograms showing the amorphous matrix co-localizing with collagen fibers. Sharp edges (traced in light peach for better visibility) delineate the amorphous matrix from empty extracellular space. Plasma membranes surrounding cells appear as strong black lines. Scale bar dimensions are annotated in the figure.

**Figure S9:**
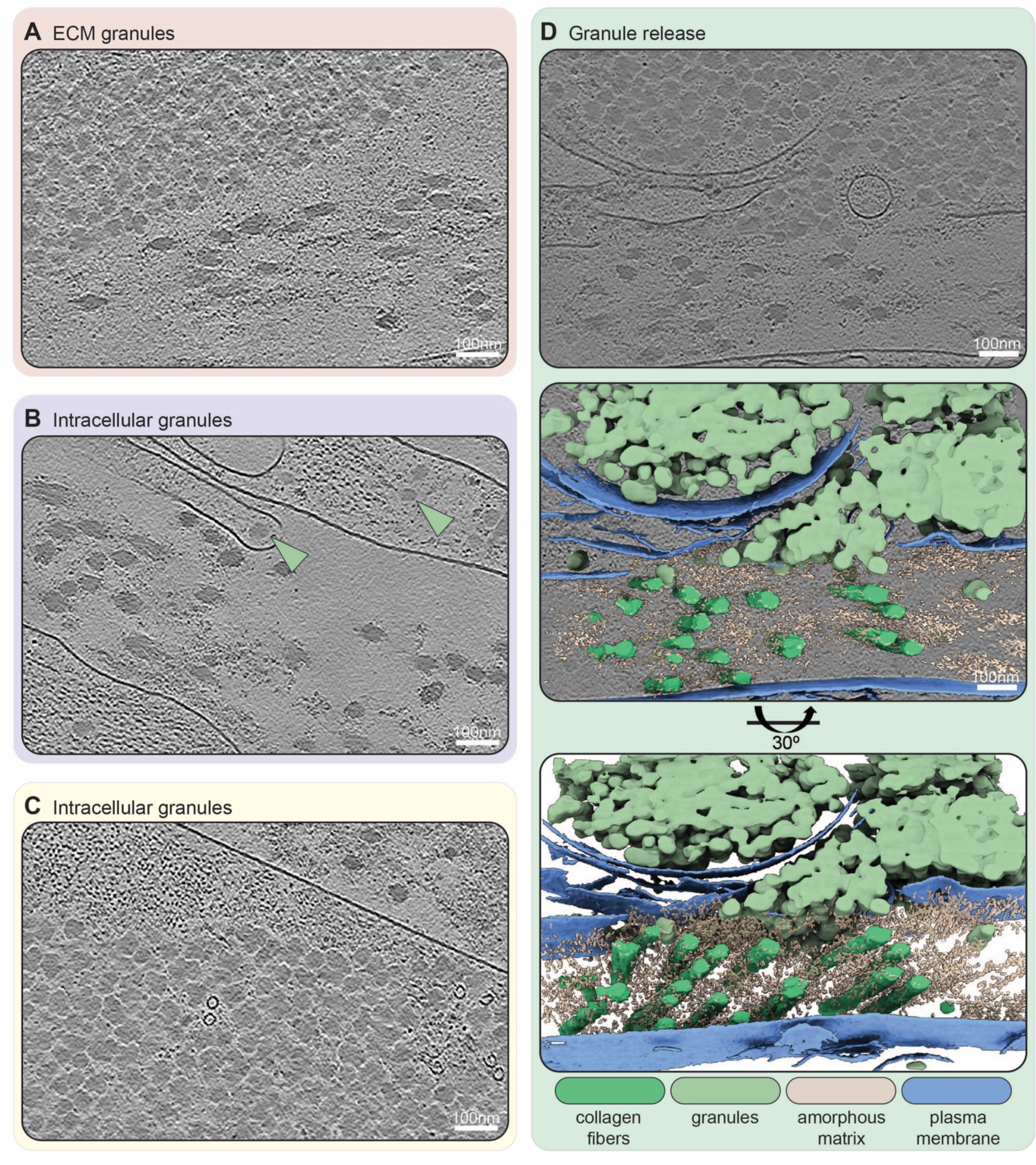
Extra- and intracellular ECM-associated granules. **A)** Granules in the extracellular space close to collagen fibers and amorphous matrix. **B)** Granules in the intracellular space, in the main cell body as well as in a cellular protrusion. Granules are indicated by a light green arrowhead. **C)** Intracellular granules close to the plasma membrane, next to actin filaments and with two microtubules interspersed. **D)** A granule release site, where granules are in both the intra- and extracellular space. The plasma membrane is partially interrupted. This tomogram was segmented using deep-learning based software (Dragonfly) and visualized using ChimeraX. The middle panel shows an overlay of the segmentation and the tomogram panel shown above. The color scheme for the different segmented cellular and ECM components is described in the figure. The lower panel shows the segmentation at an oblique angle of 30° for better visibility of the ECM fibers. All slices (1.71 nm thickness) are from IsoNet-processed tomograms. Scale bar dimensions are annotated within the tomogram.

**Table S1:**
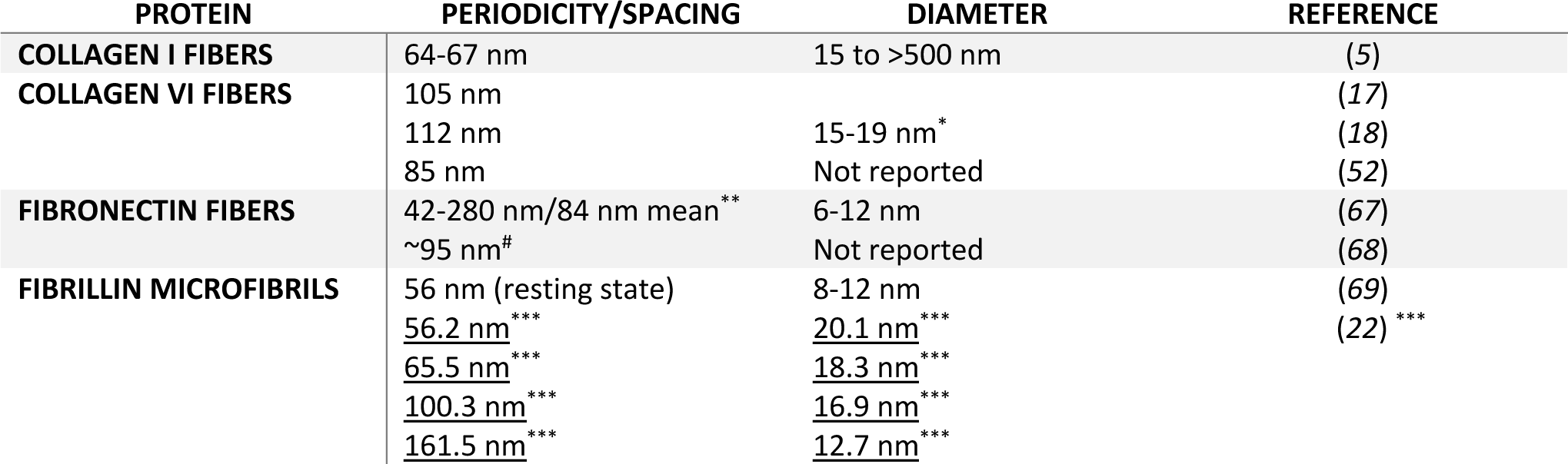
Examples of reported dimensions for different ECM fibers. Repeat patterns/periodicities/spacings and diameter measurements are given for papers when measured. We aimed to provide an overview of several studies reporting the varying measurements for the extracellular matrix filaments. We acknowledge that this table might not be complete, but still underlines the variability of filament assemblies. * double-bead region ** periodicity was reported to be diameter dependent # average epitope periodicity *** Glab and Wess (*22*) report an increased periodicity and reduced diameter following tissue extension.

**Table S2:** Predominant ECM proteins in TIFF CDMs identified by MS. TIFF CDMs were grown for 14 days and then decellularized and analyzed by LC-MS/MS. Raw data was searched against a *Homo sapiens* reference proteome and filtered for ECM proteins. ECM proteins were then sorted by the normalized log10 of their estimated expression value from highest to lowest and the top 25 hits are shown in this table. For each protein, the sequence coverage, peptide count, and spectral count are listed.

|  | PROTEIN | UNIPROT ID | SEQUENCE<br>COVERAGE | PEPTIDES<br>COUNT | SPECTRAL<br>COUNT |
| --- | --- | --- | --- | --- | --- |
| 1 | Fibronectin, Isoform 1 | P02751-1 | 76.8 | 346 | 1.155 |
| 2 | Collagen alpha-1(I) chain | P02452 | 79.8 | 268 | 408 |
| 3 | Collagen alpha-2(I) chain | P08123 | 76.4 | 179 | 277 |
| 4 | Collagen alpha-3(VI) chain | P12111 | 77.9 | 347 | 926 |
| 5 | Tenascin | P24821 | 80.9 | 190 | 520 |
| 6 | Collagen alpha-1(VI) chain | P12109 | 68.9 | 92 | 220 |
| 7 | Collagen alpha-2(VI) chain | P12110 | 53.4 | 73 | 195 |
| 8 | TGF-beta-induced protein ig-h3 | Q15582 | 73.8 | 74 | 218 |
| 9 | Galectin-1 | P09382 | 86.7 | 27 | 83 |
| 10 | Galectin-3 | P17931 | 39.2 | 15 | 38 |
| 11 | EMILIN-1 | Q9Y6C2 | 72.8 | 82 | 178 |
| 12 | Fibrillin-1 | P35555 | 62.0 | 235 | 396 |
| 13 | Serpin H1 | P50454 | 63.4 | 31 | 84 |
| 14 | Collagen alpha-1(III) chain | P02461 | 54.7 | 101 | 109 |
| 15 | Fibronectin, Isoform 11 | P02751-11 | 76 | 331 | 1.091 |
| 16 | Pyruvate kinase PKM | P14618 | 73.3 | 40 | 67 |
| 17 | Fibulin-2, Isoform 2 | P98095-2 | 49.5 | 53 | 88 |
| 18 | Collagen alpha-2(V) chain | P05997 | 39.2 | 44 | 48 |
| 19 | Collagen alpha-1(V) chain | P20908 | 29.4 | 47 | 62 |
| 20 | Collagen alpha-1(XII) chain | D6RGG3 | 65.2 | 190 | 277 |
| 21 | Tenascin, Isoform 4 | P24821-4 | 82.7 | 186 | 516 |
| 22 | Decorin | P07585 | 74.1 | 29 | 48 |
| 23 | BM-specific heparan sulfate<br>proteoglycan core protein | P98160 | 66.0 | 214 | 357 |
| 24 | Fibulin-1 | P23142 | 68.4 | 39 | 72 |
| 25 | Fibrillin-2 | P35556 | 48.5 | 136 | 201 |

**Table S3:** Scouting of different cryoprotectants and buffers for their vitrification potential. An overview table showing the tested cryoprotectant/buffer combinations, detailing the achieved vitrification status and the introduced background for each combination. Vitrification was judged according to the occurrence of reflections caused by hexagonal ice crystals in the cryo-lift out lamellae. Vitrification was judged as *complete* if no reflections could be detected and as *incomplete* if reflections of any severity were detected. Background was judged as *none* if no cryoprotectants were added, as *acceptable* if structures within the cryo-lift out lamellae were still distinctively visible, and as *high* if the introduced background obscured structures within the cryo-lift out lamellae.

| Cryoprotectant | Degassed | Vitrification | Background |
| --- | --- | --- | --- |
| Native | - | Incomplete | None |
| 15% PVP in PBS | - | Incomplete | Acceptable |
| 15% PVP in 0.1 M PB | - | Incomplete | Acceptable |
| 15% BSA in 0.1M PB | - | Incomplete | High |
| 20% Dextran/5% Sucrose in PBS | - | Complete | High |
| 20% Dextran/5% Sucrose in PBS | + | Complete | High |
| 20% Dextran in PBS | + | Incomplete | Acceptable |
| 10% Dextran in PBS | + | Incomplete | Acceptable |
| 10% Dextran in PB | + | Complete | Acceptable |
| 5% Dextran in PB | + | Incomplete | Acceptable |
| Native | + | Incomplete | None |
| 10% BSA in medium | + | Incomplete | Acceptable |
| 20% BSA in medium | + | Complete | High |

## Supplementary Movies

**Movie S1:** Movie of the tomogram and segmentation shown in Figure 3A.

**Movie S2:** Movie of the tomogram and segmentation shown in Figure 3B.

**Movie S3:** Movie of the tomogram and segmentation show in Figure S9D.

**Movie S4:** Movie showing features following a linear filament-like trajectory, annotated with a black arrowhead. The small bead-like filaments are annotated with yellow arrowheads or a yellow ellipsoid.

**Movie S5:** Movie showing features following a linear filament-like trajectory, annotated with a black arrowhead.

